# Amplified genome editing by *in vivo* editor production

**DOI:** 10.64898/2026.01.13.699115

**Authors:** Wayne Ngo, Daniel Rosas-Rivera, Kevin M. Wasko, Longhui Qiu, Min Hyung Kang, Shaan Gogna, Jingkun Zeng, Mason T. Hooks, Jamie L.Y. Wu, Zhongmei Li, Jennifer A. Doudna

**Affiliations:** Innovative Genomics Institute; University of California, Berkeley; Berkeley CA, USA; Gladstone Institute of Data Science and Biotechnology; San Francisco, CA, USA; California Institute for Quantitative Biosciences, University of California, Berkeley; Berkeley, CA, USA; Department of Molecular and Cell Biology, University of California, Berkeley; Berkeley, CA, USA; Department of Medicine, University of California, San Francisco; San Francisco, CA, USA; Gladstone-UCSF Institute of Genomic Immunology; San Francisco, CA, USA; Howard Hughes Medical Institute, University of California, Berkeley; Berkeley CA, USA; Molecular Biophysics and Integrated Bioimaging Division, Lawrence Berkeley National Laboratory; Berkeley, CA, USA; Department of Chemistry, University of California, Berkeley; Berkeley, CA, USA; Li Ka Shing Center for Genomic Engineering, University of California, Berkeley, Berkeley, CA, USA

## Abstract

Genome editing enzymes have vast therapeutic potential. However, achieving sufficient delivery *in vivo* remains a major challenge, because editing machinery is confined to the subset of transfectable cells in a tissue. Here, we tested the possibility that genome editing could be amplified *in vivo* by enabling transfected cells to transfer editing enzymes to neighboring cells. Our data show that this NANoparticle-Induced Transfer of Enzyme (NANITE) strategy quadrupled editing efficiency in cultured cells relative to non-spreading controls. A single intravenous injection of the NANITE plasmid into mice induced ∼3-fold higher levels of liver editing at the transthyretin (*Ttr*) locus relative to non-spreading controls, with corresponding reductions in serum TTR levels. Spreading therapeutic enzymes offers a nonviral and non-infectious strategy to boost therapeutic effects after delivery.

## Main

Genome editing enzymes promise to revolutionize medicine by overriding or correcting the underlying genetic basis of disease. Editing enzymes are delivered as ribonucleoproteins (RNPs) or their encoding nucleic acids in protective delivery vehicles that circulate around the body and enter target cells. Treatment requires editing a sufficient portion of target cells to elicit a therapeutic effect^1^. Researchers have focused on surpassing these thresholds by engineering delivery vehicles to transfect more cells. Beyond engineering delivery vehicles, strategies to increase the number of edited cells after delivery could also increase therapeutic effects. Two approaches have been described to increase the portion of edited cells after delivery. First, edited cells can be stimulated to undergo clonal expansion. Antigen-dependent expansion was used to increase genome-edited T cell numbers *in vivo*. Although only a small number of T cells were initially transfected and edited, exposure to antigens drove expansion until edited T cells were up to 20% of all T cells^2^. Second, edited cells can repopulate a tissue after depletion of unedited cells. Repair Drive uses a siRNA against an essential hepatocyte gene (Fah) to selectively kill unedited cells while edited cells express an siRNA-resistant Fah variant and expand to repopulate the liver.^3^ A similar method using antibody-depletion has been described for hematopoietic stem cells.^4^ These approaches rely on cell-specific clonal expansion or deliberate killing of unedited cells, which limits their generalizability and safety.

We wondered if it is possible to directly edit more cells after delivery without clonal expansion or killing unedited cells. Editing is currently confined to transfected cells, because large hydrophilic RNPs cannot transfer from transfected to untransfected cells after delivery. Intercellular transfer of proteins and RNA can occur endogenously through exosomes^5^ and tunneling nanotubes^6^. Engineered systems have achieved transfer of proteins or RNA *in vitro*^7–10^ and *in vivo*^11–15^ using vesicles. However, it is unclear whether the intercellular transfer of protein-RNA complexes, such as CRISPR-Cas9 RNPs, after delivery could increase genome editing *in vivo*. Here, we explored whether genome editing efficiencies could be increased by enabling transfected cells to transfer Cas9 RNPs to surrounding cells. We hypothesized that enabling cells to transiently produce enveloped delivery vehicles (EDVs) *in situ* could spread editing enzymes and increase genome editing after delivery (Fig. 1a). EDVs are virally-derived lipid vesicles that package *Streptococcus pyogenes* Cas9 (SpCas9) RNPs^16–18^. We previously used EDVs as standalone delivery vehicles that were produced and purified before administration to recipient cells *in vitro* and *in vivo* for genome editing^16–18^. In this study, we found that delivering plasmids encoding EDV components to cells *in vitro* and *in vivo* amplified genome editing three-to four-fold relative to non-spreading controls without toxicity. This NANoparticle Induced Transfer of Enzymes (NANITE) strategy amplifies editing without clonal expansion and killing unedited cells, offering a generalizable method to enhance therapeutic genome editing.

**Figure 1.**
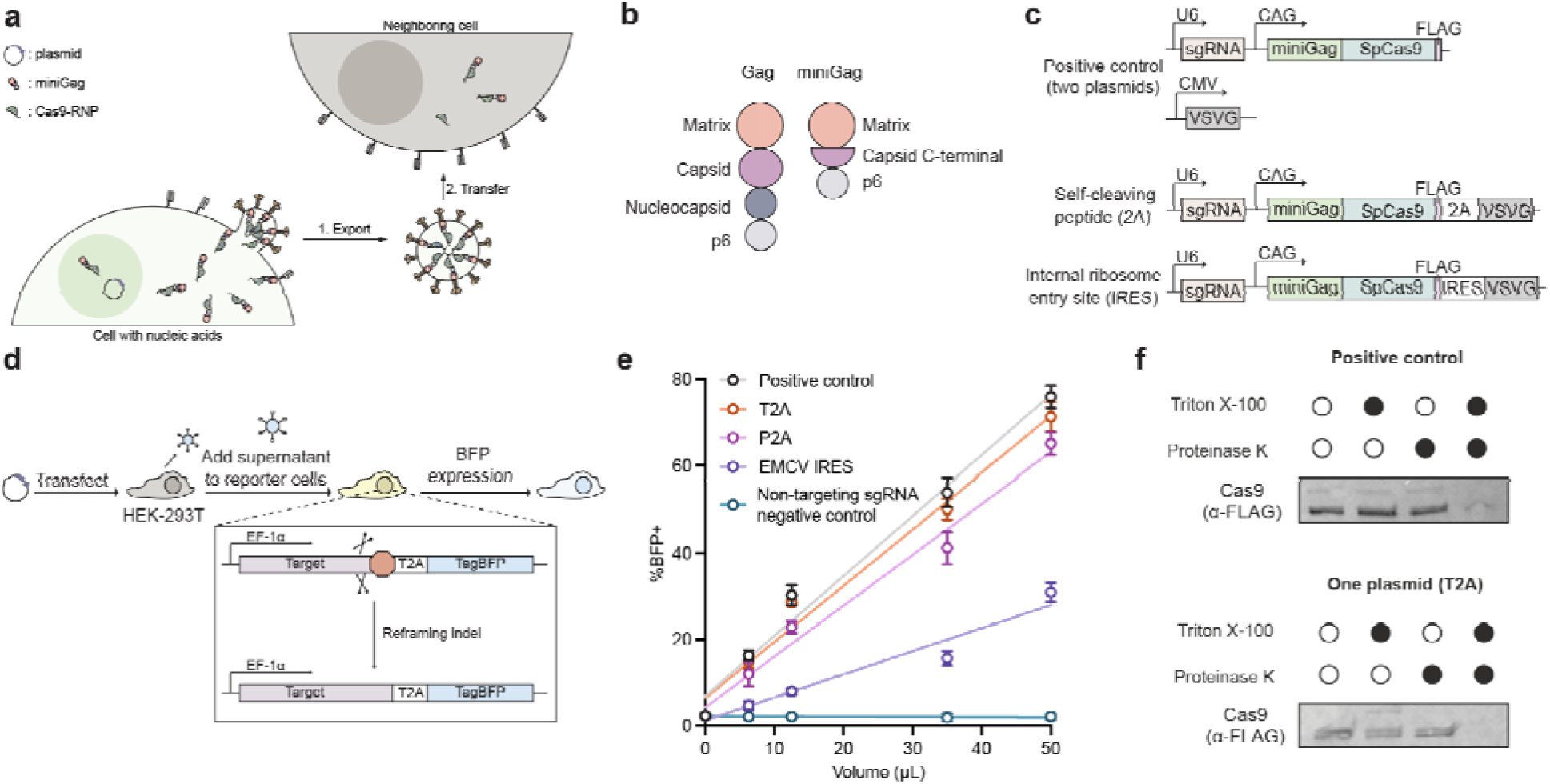
Genome editing vesicles can be produced from a single plasmid. **(a)** Schematic of NANITE, in which cells that receive nucleic acid cargo transiently produce and secrete Cas9 RNPs in vesicles. Locally produced vesicles subsequently transduce neighboring cells to spread genome editing. **(b)** Schematic of the miniGag structural protein compared to HIV-1 Gag structural protein. **(c)** Design of NANITE. The positive control consists of the two-plasmid system to produce miniEDVs, in which one plasmid encodes the cargo and internal structural components of the vesicle and a second plasmid encodes the surface protein necessary for cell transduction. In NANITE, the surface protein is encoded on the first plasmid, separated by either self-cleaving peptides or internal ribosome entry sites. **(d)** Experimental scheme for testing NANITE-plasmids compared to the positive control. HEK-293T producer cells were transfected with plasmids. Supernatant containing vesicles was harvested 48 hours post-transfection and incubated with reporter cells that express blue fluorescent protein (BFP) upon genome editing. Schematics (a–d) are not to scale. **(e)** Fraction of reporter cells expressing BFP after incubation with supernatants quantified by flow cytometry. Separating the surface protein from internal components using a T2A sequence resulted in vesicles with comparable activity to the positive control. The non-targeting sgRNA negative control is a miniEDV with a non-targeting gRNA that does not restore BFP expression. Data are presented as mean ± SD of three replicates. **(f)** Protease protection assay of vesicles produced by either the positive control or T2A design. A FLAG tag is present on the C-terminus of Cas9, allowing internal components of the vesicle to be detected by anti-FLAG antibodies. Cas9 is enclosed within vesicles in both conditions, as both Triton X-100 and proteinase K are required for degradation. The assay was repeated twic with similar results.

## Results

### Encoding NANITE on a single plasmid

We previously showed that fusing SpCas9 RNPs to truncated lentiviral Gag structural proteins (“miniGag”, Fig. 1b) allows cells to produce EDVs that encapsulate editing enzymes^18^. These vesicles are non-replicative because they do not package nucleic acids encoding their components. Cells receiving these vesicles will be edited but unable to produce more vesicles^18^. We wanted to explore whether transient *in situ* production of EDVs could amplify genome editing effects. To explore this possibility, we first addressed the complexity of production which required co-delivery of multiple plasmids at fixed ratios which is difficult to control *in vivo*^16–20^. We designed single plasmids to express all the components to assemble and secrete SpCas9 RNP-containing EDVs (Fig. 1c). Single-guide RNAs (sgRNAs) were encoded on the same plasmid and expressed from a U6 promoter. To test for EDV production, plasmids were transfected into HEK-293T cells, and the supernatant was harvested and incubated with reporter cells that express BFP upon genome editing (Fig. 1d and Supp. Fig. 1a-c). All single-plasmid constructs produced genome editing-competent EDVs as detected by flow cytometry (Fig. 1e, flow gating shown in Supp. Fig. 1d, e). The T2A design performed nearly identically to the two-plasmid positive control, while the P2A and encephalomyocarditis virus internal ribosomal entry site (IRES) designs showed reduced activity. Western blots of the cell lysates confirmed complete T2A-catalyzed post-translational cleavage of the Vesicular Stomatitis Virus Glycoprotein (VSVG) fusogen, with miniGag-Cas9 and VSVG detected as separate bands (Extended Data Fig. 1a). Western blots of vesicles revealed slightly reduced VSVG loading on the vesicles, consistent with VSVG expression relying on a T2A self-cleaving sequence rather than an independent promoter (Extended Data Fig. 1b). The T2A design had a similar rate of vesicle production, Cas9 loading, and sgRNA loading to the two-plasmid positive control (Extended Data Fig. 1c-e). A protease protection assay confirmed that Cas9 RNPs were packaged inside vesicles, because internal components were protected from proteinase K degradation unless Triton X-100 disrupted the membrane (Fig. 1f). These results demonstrate that a single plasmid can encode fully functional EDVs. We used the T2A design for subsequent experiments, labelled as ‘NANITE’.

### NANITE amplifies editing in situ

We next explored whether *in situ* production of EDVs would allow editing enzymes to spread from transfected to untransfected cells. We added an IRES-zsGreen cassette to the NANITE plasmid for labeling transfected cells and designed a control plasmid expressing only Cas9 RNP and zsGreen (Fig. 2a). Plasmids were complexed with polyethylenimine and delivered into BFP reporter cells^21,22^. Transfected (zsGreen+) and edited (BFP+) cells were quantified by flow cytometry 48 hours after transfection (Fig. 2b, gating in Supp. Fig. 1d). Transfecting cells with the Cas9 RNP-plasmid yielded 32% transfected (zsGreen+) and 19% edited (BFP+) cells. All edited cells were transfected, but only a fraction of the transfected cells were edited due to insufficient Cas9 RNP production or indels that did not restore BFP expression (Supp. Fig. 1a). Transfecting cells with NANITE-plasmids produced 34% transfected and 57% edited cells. Notably, 33% of cells were edited without transfection (BFP+, zsGreen-) (Fig. 2b). Across transfection efficiencies, only ∼87% of cells transfected with Cas9 RNP-plasmids became edited, while the fraction of cells edited with NANITE-plasmids averaged 293% of the transfected fraction with a maximum of 386% (Fig. 2c). To confirm that the multicistronic NANITE-plasmid design did not reduce zsGreen expression and underestimate transfected cells, we inserted an inert stuffer into the Cas9 RNP-plasmid to match the promoter-to-IRES distance in NANITE (Supp. Fig. 2a). These plasmids behaved similarly to the Cas9 RNP-plasmid, consistent with IRES-driven translation being insensitive to distance from the cap of the mRNA (Supp. Fig. 2b). These results show that NANITE-plasmids edit more cells than are initially transfected.

**Figure 2.**
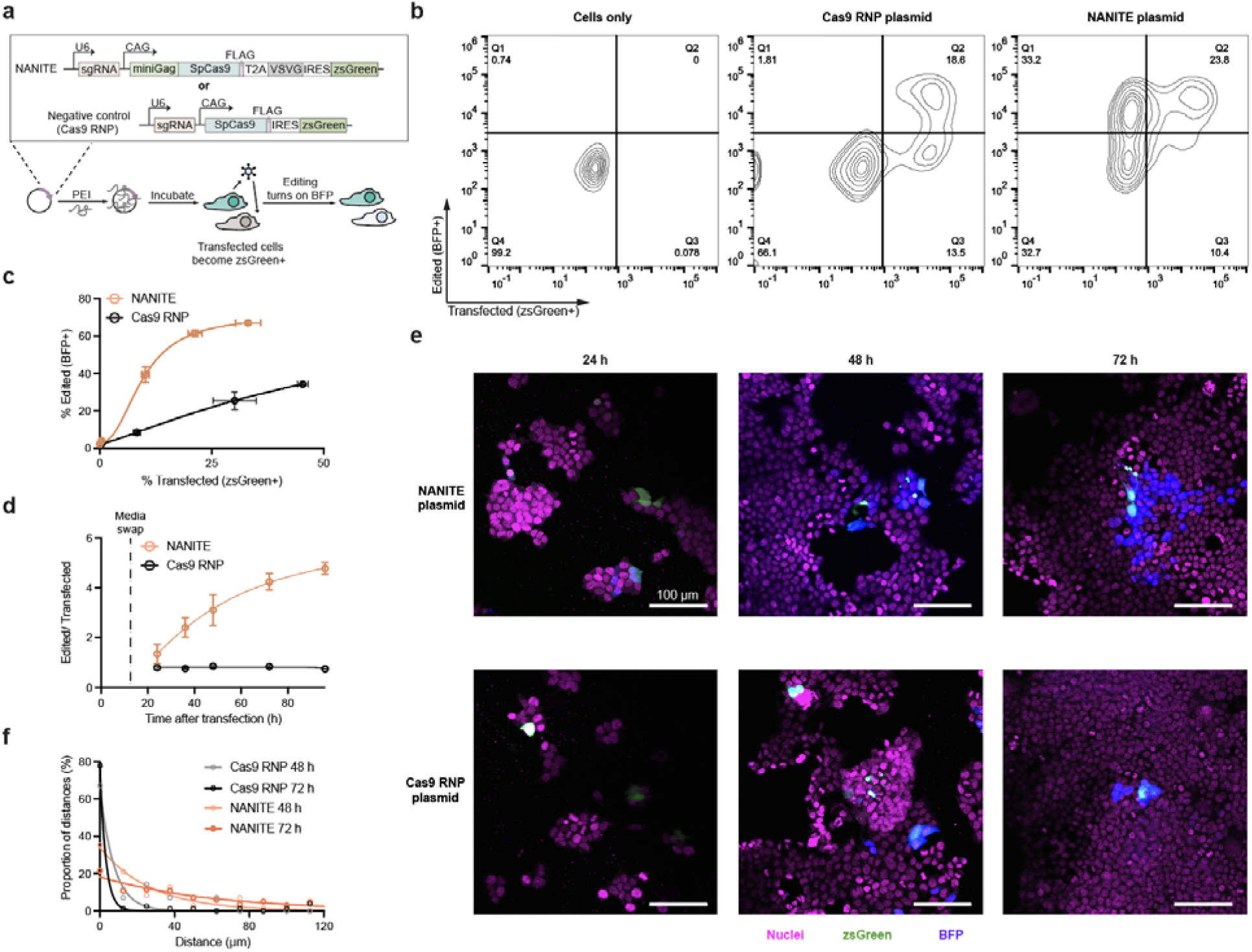
NANITE spreads genome editing *in situ*. **(a)** Experimental scheme for testing NANITE spread. Plasmids were complexed with polyethylenimine and incubated with reporter HEK-293T cells. Transfected cells express zsGreen. Genome editing turns on BFP expression. Schematic is not to scale. **(b)** Representative flow cytometry plots showing zsGreen and BFP expression. Transfection with NANITE-plasmids resulted in a population of cells that were edited (BFP+) but not transfected (zsGreen-). **(c)** Fraction of edited cells relative to transfected cells following transfection with NANITE- or Cas9 RNP-plasmids. More edited cells than transfected cells were observed in the NANITE-plasmid condition. Data are presented as mean ± SD of three replicates. **(d)** Ratio of edited to transfected cells over time after transfection with NANITE- or Cas9 RNP-plasmids. NANITE-transfected cells accumulated 4.3 ± 0.3 edited cell per transfected cell over 72 hours, compared to 0.8 edited cells per transfected cell in Cas9 RNP-plasmid controls. Data are presented as mean ± SD of three biological replicates. **(e)** Representative confocal microscopy images of cells transfected with NANITE- or Cas9 RNP-plasmids. NANITE-transfected cells edit neighboring cells. Scale bar as indicated. Two biological replicates were performed. **(f)** Distribution of distances between edited (BFP+) and nearest transfected (zsGreen-) cells, fitted to an exponential decay. Six fields of view from two biological replicates were analyzed. NANITE increases the distance between edited cells and their nearest transfected cell, indicating spread of genome editing.

We tracked the fraction of edited and transfected cells over time to quantify how NANITE spreads editing. In NANITE conditions, each transfected cell edited one additional cell every 12 hours, plateauing at 4.3 ± 0.3 edited cells per transfected cell after 72 hours (Fig. 2d). Cells transfected with Cas9 RNP plasmids remained stable at ∼0.8 edited cells per transfected cell across all timepoints. To visualize how NANITE was spreading, we imaged cells after low transfection to observe discrete spreading events (Fig. 2e). Transfected zsGreen+ cells appeared within 24 hours in both Cas9 and NANITE conditions. BFP+ edited cells emerged after 48 hours. We observed clusters of edited cells surrounding cells transfected with NANITE. We measured distances from each edited cell to the nearest transfected cell center-to-center using Cellprofiler^23^. These distances were binned into histograms, then fitted to a one-phase exponential decay (Fig. 2f). Cas9 RNP plasmid half-distances were 2-5 μm at 48 and 72 hours, which was less than the average cell radius (12 ± 3 μm), indicating no spreading. NANITE plasmid half-distances were 19 and 42 μm at 48 and 72 hours respectively, showing progressive radial spread of editing enzymes multiple cell lengths from initially transfected cells.

To determine if secreting Cas9 decreased the concentration of Cas9 inside the cell over time, we isolated transfected cells using fluorescence activated cell sorting, then quantified Cas9 using enzyme-linked immunosorbent assays (ELISA) over time (Extended Data Fig. 2a). Cas9 concentrations declined over four days despite stable cell numbers, suggesting that Cas9 was exported from the cell and that transient NANITE expression did not kill producer cells (Extended Data Fig. 2b, c).

We then tested how NANITE was spreading. We first separated NANITE-transfected cells from receiver cells using a 3.0 µm membrane in transwell assays to block direct cell contact, which is required for cell–cell fusion,^24^ trogocytosis,^25,26^ and tunneling nanotube-mediated transfer^27^. We found that cells were edited in the bottom well even though transfected cells were seeded only in the top well. Editing increased with the number of NANITE-transfected cells in the top well (Extended Data Fig. 3a, b). Results were similar when the seeding orientation was reversed (Extended Data Fig. 3c). No editing was detected in the opposing well when cells were transfected with Cas9 RNP-plasmids in either orientation, suggesting no uptake of free Cas9 RNP. These data argue that editing was not spread through direct contact-dependent mechanisms. We then used small-molecule inhibitors of different transfer pathways to further isolate the mechanism of spread. Inhibitors were added after transfection to avoid confounding effects on transfection efficiency (Extended Data Fig. 3d). DMSO did not affect spread and Cas9 RNP plasmid-transfected cells did not induce editing in receiver cells (Extended Data Fig. 3e). Inhibitors of tunneling nanotube formation (L778123)^28^ and PI3K-dependent membrane reorganization (LY294002)^29^ also did not reduce editing in receiver cells at non-cytotoxic concentrations (Extended Data Fig. 3f). Similarly, inhibitors of microvesicle (Y27632)^30^ and exosome (GW4869)^31^ biogenesis did not impair spread, suggesting that spread was occurring through EDVs and not endogenous exosomes. In contrast, inhibitors of endosomal acidification (bafilomycin A1)^32^ and microtubule-dependent endocytic uptake (colchicine)^33^ significantly reduced editing in receiver cells. Together, our data argue that NANITE spreads in an endocytosis-dependent manner consistent with VSVG-mediated transfer, although we cannot rule out that other mechanisms may contribute in different cellular contexts.

We wondered whether NANITE functions in other cell types. We compared editing mediated by NANITE-or Cas9 RNP-plasmids at the endogenous *B2M* locus in HepG2 liver cells, which are commonly used as an *in vitro* model for hepatocytes^34^. Edited cells in the NANITE-plasmid condition averaged 238% of the transfected cells compared to ∼92% in the Cas9-plasmid condition (Extended Data Fig. 4a). Similar results were observed in C2C12 muscle cells, where edited cells in the NANITE-plasmid condition averaged 387% of the transfected population (Extended Data Fig. 4b). Unlike approaches that depend on clonal expansion or depletion of unedited cells, our data shows that NANITE can be applied to multiple cell types without additional engineering.

### The tropism of NANITE can be tuned

Since NANITE functions in multiple cell types, we sought to control its tropism. Previous studies showed that EDV tropism can be tuned by presenting single-chain antibody fragments (scFvs) with receptor-binding deficient fusogens on the vesicle surface^16,35,36^. We explored whether NANITE tropism could be similarly controlled by designing NANITE-encoded vesicles to display both scFvs and receptor-binding deficient VSVG (VSVGmut) simultaneously (Fig. 3a). CD19-and ACE2-binding scFvs were selected because they were validated and their receptors were not endogenously expressed in HEK-293T cells, enabling specificity testing without confounding background receptor expression^37,38^. We transfected HEK-293T cells with targeted NANITE-plasmids and harvested supernatants for incubation with recipient cells either expressing or lacking the cognate receptor (Fig. 3b). Editing at the *B2M* locus was quantified by flow cytometry (Supp. Fig. 3a, b). VSVG-NANITE edited all cell types equally (Fig. 3c). VSVGmut-NANITE showed background editing, especially at high doses, consistent with the presence of non-receptor-specific uptake pathways in HEK-293T cells^39^ and prior reports of low-level nonspecific uptake of VSVGmut particles^36,40^. Anti-CD19-NANITE and Anti-ACE2-NANITE showed up to 25-fold and 11-fold more editing of cells expressing their matching receptors, demonstrating that NANITE tropism can be engineered. We next wondered how tropism functions with *in situ* production of NANITE-encoded vesicles. We transfected mCherry+ producer cells and co-cultured them at a 1:1:1 ratio with CD19-expressing (mNeonGreen+) and ACE2-expressing HEK-293T receiver cells (Fig. 3d, flow cytometry gating shown in Supp. Fig. 3c). The transfection efficiencies of the producer cells are shown in

**Figure 3.**
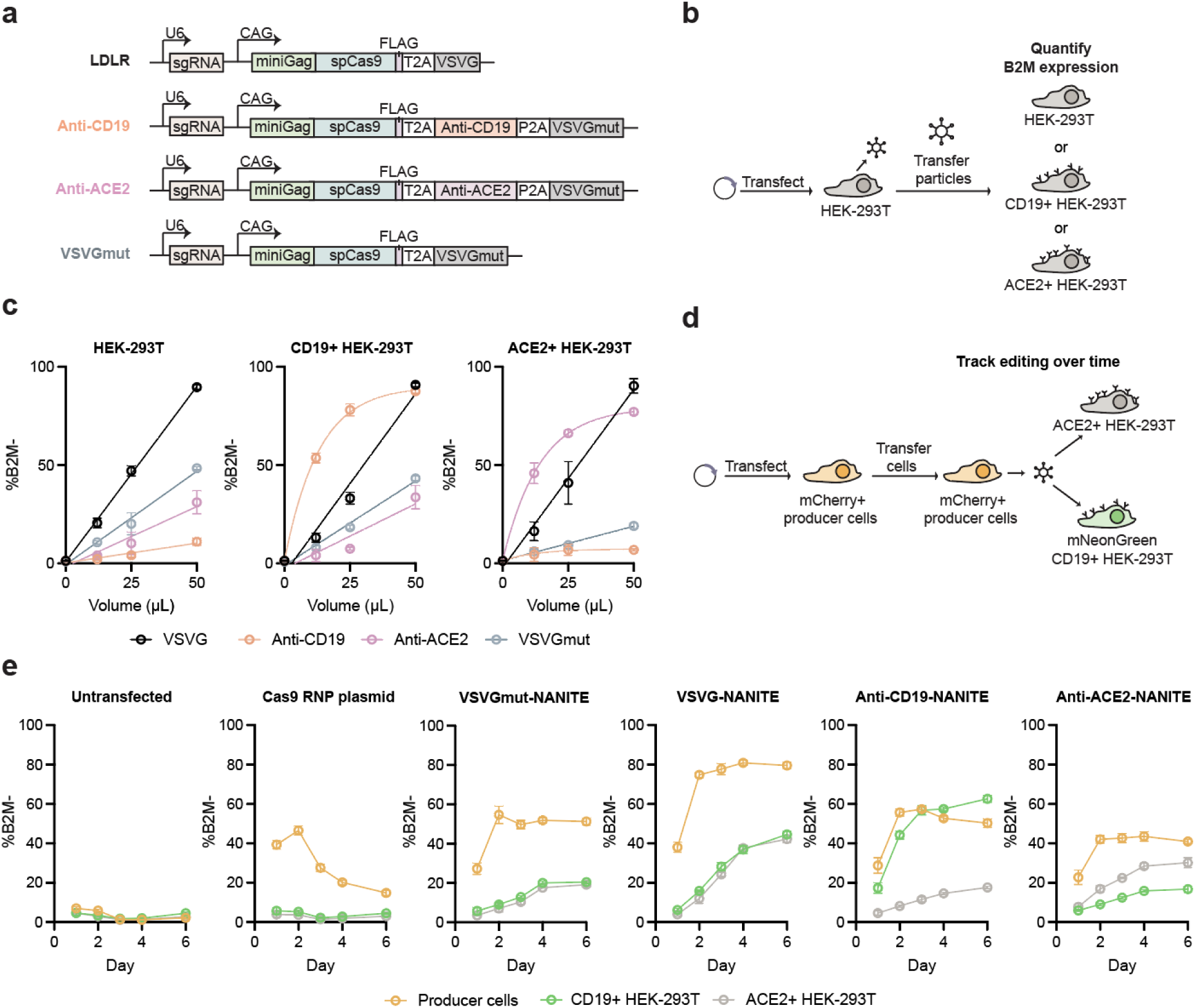
NANITE tropism can be controlled. **(a)** Schematics of targeted NANITE designs. Single-chain antibody fragments are displayed with VSVGmut. **(b)** Experimental scheme for testing targeted NANITE designs. HEK-293T cells were transfected, vesicles were harvested, and then added to recipient cells. Schematics are not to scale. **(c)** Fraction of edited HEK-293T, CD19+ HEK-293T, or ACE2+ HEK-293T cells after incubation with VSVG-(black), VSVGmut- (gray), anti-CD19- (orange), or anti-ACE2- (pink) NANITE-encoded vesicles. Targeted NANITE-encoded vesicles were specific for cells expressing the cognate receptor. Data are presented as mean ± SD of three replicates. **(d)** Experimental scheme for testing targeted NANITE-plasmids *in situ*. Producer cells (mCherry+) were transfected with plasmids and co-cultured in a 1:1:1 mixture with CD19-expressing HEK-293T (mNeonGreen+) and ACE2-expressing HEK-293T cells. Editing at the B2M locus was quantified by flow cytometry at different time points. **(e)** Fraction of edited (B2M-) producer and receiver cells over time. Targeted NANITE-plasmids showed tropism for their cognate cells. Data presented as the mean ± SD of three biological replicates.

Supplementary Figure 4. No editing was observed in the absence of transfection (Fig. 3e). Cas9 RNP-transfected producer cells showed editing only in producer cells, with no detectable transfer to either receiver population over 6 days. VSVGmut-NANITE produced low-level editing in both receiver cell types, reaching ∼20% by day 6. VSVG-NANITE edited up to 46% of receiver cells without discrimination between populations. Anti-CD19-NANITE showed preferential transfer to CD19-expressing cells (∼60% edited) relative to ACE2-expressing cells (∼18% edited). Conversely, Anti-ACE2-NANITE showed preferential transfer to ACE2-expressing cells over CD19-expressing cells. Together, these results demonstrate that the tropism of NANITE can be directed by engineering the vesicle surface.

Beyond controlling tropism using scFvs, we reasoned that cell-type specificity could also be controlled using tissue-specific promoters. We replaced the CAG promoter with liver-specific TGB610, GFAP and LP1 promoters (Extended Data Fig. 5a) and transfected equimolar amounts into HEK-293T or HepG2 cells to assess NANITE activity as measured by BFP expression^41–43^ (Extended Data Fig. 5b). To compare the promoters across cell lines, NANITE activity was normalized to the CAG promoter within that cell type. The TGB610 promoter showed significantly higher activity in on-target HepG2 compared to off-target HEK293T cells, but with only 34% the activity of CAG (Extended Data Fig. 5c). Prior studies have shown a trade-off between promoter expression strength and specificity^44^. Nevertheless, these results show the possibility of controlling NANITE expression through promoter engineering, allowing us to control the tropism of NANITE at multiple stages of deployment.

### NANITE amplifies in vivo genome editing efficiency

Having shown that NANITE can amplify genome editing efficiency in cultured cells by disseminating editing enzymes, we next explored whether NANITE amplifies *in vivo* genome editing efficiency. We chose to test NANITE for transthyretin (TTR) liver editing in mice, because TTR is an established therapeutic target for transthyretin amyloidosis with validated sgRNAs and characterization assays^45,46^. We used the NANITE design with the CAG promoter and VSVG, as CAG showed the highest *in vitro* activity and VSVG naturally transduces hepatocytes expressing the low-density lipoprotein receptor (LDLR). NANITE plasmids were formulated for *in vivo* delivery using lipid nanoparticles (LNPs, ∼109 nm in diameter with ∼85% encapsulation efficiency) containing a cGAS-STING inhibitor (Supp. Fig. 5a, b)^47^. Although these NANITE-LNPs showed amplified editing *in vitro* (Supp. Fig. 5c), *in vivo* doses up to 25 µg (matching the highest dose previously reported^47^) produced no detectable transfection (Supp. Fig. 5d, e). This prompted us to explore alternative *in vivo* plasmid delivery methods.

We subsequently turned to hydrodynamic injection, a validated method for transiently delivering plasmids to the liver that has been explored in humans for liver gene therapy^48–50^. Dose optimization showed that 20 pmol of plasmid injected hydrodynamically could transfect ∼5% of liver cells (Supp. Fig. 6), consistent with published efficiencies for 10-15 kb plasmids^51^. To test NANITE *in vivo,* we injected 20 pmol of either NANITE- or Cas9 RNP-expressing plasmids into mice at 20 pmol (Fig. 4a). Similar transfection efficiencies were observed between NANITE- and Cas9 RNP- at 24 hours, as assessed by zsGreen fluorescence imaging (8 ± 3% and 6 ± 2%, respectively; Fig. 4b, c) and flow cytometry (Supp. Fig. 7). We quantified transfection at 24 hours, because plasmid expression is known to be maximal at 24 hours in mice^52,53^. Western blot analyses of liver lysates confirmed NANITE protein production *in vivo*, albeit at lower levels than Cas9 protein production, likely due to NANITE’s larger size (Extended Data Fig. 6a). A similar fraction of cells were transfected with both plasmid conditions with higher protein expression in the Cas9 RNP-plasmid group.

**Figure 4.**
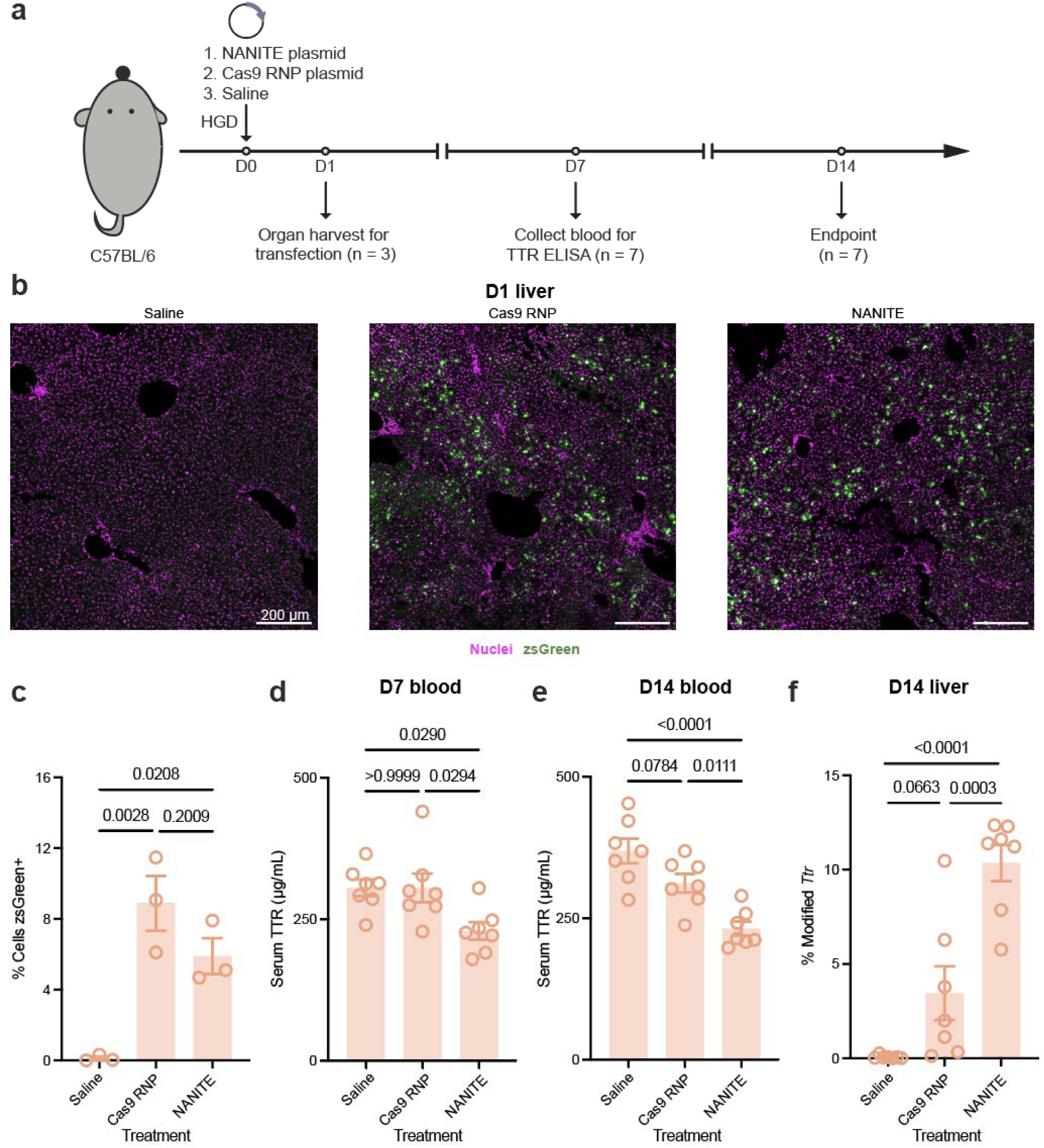
NANITE amplifies genome editing *in vivo.* **(a)** Experimental scheme for genome editing *Ttr* in mice. Mice were hydrodynamically injected (HGD) with plasmids. Organs were collected 24 hours post-injection to quantify transfection efficiency (n = 3). Editing was monitored by TTR ELISA on blood collected on days 7, and 14. Tissues were harvested at day 14 to assess editing and animal health (n = 7). **(b)** Representative immunofluorescence images of liver sections showing nuclei (magenta) and zsGreen (green). Scale bar as indicated. **(c)** Fraction of transfected cells in the liver. One field of view per mouse was quantified. No significant difference in the fraction of transfected cells was observed between NANITE- and Cas9 RNP-plasmid conditions. **(d)** Serum TTR concentration 7 days post-administration. **(e)** Serum TTR concentration 14 days post-administration. NANITE significantly reduced TTR levels. **(f)** Next-generation sequencing of liver genomic DNA harvested 14 days post-injection. NANITE-plasmid treated mice showed greater editing at the *Ttr* locus compared to saline or Cas9 RNP-plasmid treated mice. For (c–f), statistical significance was determined by one-way ANOVA with Tukey’s multiple comparison test. Relevant p-values are indicated. Data are presented as mean ± SEM (c: n = 3 mice; d–f: n = 7 mice).

Since TTR is secreted into the bloodstream, we monitored editing non-invasively by performing quantifying serum TTR concentrations with ELISAs seven and 14 days post-administration to allow for NANITE or Cas9 RNP expression, *Ttr* editing, and equilibration of serum TTR levels. NANITE-treated mice showed significantly reduced serum TTR levels (Fig. 4d), persisting to day 14 when NANITE mice had 60 ± 11 % the TTR concentration of saline controls (Fig. 4e). In contrast, Cas9 RNP plasmid-treated mice did not have statistically significant reductions in TTR (85 ± 17%). Since serum TTR levels plateaued between days 7 and 14, we terminated the experiment at day 14. At the endpoint, next-generation sequencing of DNA extracted from liver tissue revealed 11 ± 3 % TTR editing in the NANITE-treated mice, versus 4 ± 4 % TTR editing in the Cas9-RNP-treated mice (Fig. 4f). Thus, NANITE outperformed Cas9 RNP despite the Cas9 RNP-plasmid showing higher protein expression at comparable transfection levels (Extended Data Fig. 6a). To contextualize the level of editing, we compared the observed extent of NANITE-mediated genome editing to that observed using miniEDVs for *in vivo* delivery^18^. The 20 pmol dose of NANITE plasmid achieved editing comparable to 5.5 × 10^10^ miniEDVs (1.5 × 10^12^ sgRNAs) per mouse produced from 49 pmol of total plasmid (∼120 million producer cells) (Supp. Fig. 8a - d). Together, these results are consistent with NANITE amplifying the number of cells that are edited after a single injection *in vivo*, relative to non-amplifying delivery modalities.

To examine the safety of NANITE treatment, we first examined the persistence of the plasmids by performing qPCR on mouse liver DNA extracted on day one and 14 post-injection. By day 14, plasmids were undetectable in two of the Cas9 RNP-treated mice and one of the NANITE-treated mice, with mean copy numbers of 1.7 ± 1.6 and 4.7 ± 4.9 per cell, respectively. The near-complete loss of plasmid signal indicates that the plasmids are not stably maintained (Extended Data Fig. 6b - d). Immunohistochemistry and Western blotting detected no zsGreen or FLAG-tagged protein 14 days post-administration (Extended Data Fig. 6e - g), confirming transient expression. These findings demonstrate that NANITE expression is transient *in vivo*.

No significant off-target editing was detected at ten predicted loci (Extended Data Fig. 7a - k). RT-qPCR also did not detect elevated sgRNAs concentrations in serum from NANITE-treated mice compared to Cas9 RNP or saline controls (Extended Data Fig. 7l, m), confirming that NANITE activity was confined to the liver. Concomitantly, no editing occurred in ovaries (Extended Data Fig. 7n). All mice experienced an initial 5-10% weight loss post-injection, recovering by day four, with no group differences (Extended Data Fig. 8a). Blinded histopathology scoring on the liver did not show increased steatosis, inflammation, fibrosis, mitosis, glycogen accumulation, necrosis, karyomegaly, bile duct inflammation, mineralization, or fused multinucleated cells in NANITE- or Cas9 RNP-plasmid-treated mice (Extended Data Fig. 8 b-n). Importantly, we did not detect increased inflammation, immune cell infiltration, necrosis, or fibrosis which would be expected if transfected cells were eliminated. There was no evidence of liver cell proliferation, as mitosis scores were not significantly increased (Extended Data Fig. 8h). To evaluate physiological functions of the liver and other organs, we examined blood biochemistry. Aspartate aminotransferase (AST) and alanine aminotransferase (ALT) levels in NANITE plasmid-treated mice were comparable to Cas9 RNP plasmid-treated mice, and moderately elevated compared to saline controls on day seven. The elevation was attributable to plasmid treatment rather than NANITE specifically. All values remained within the normal physiological range (Extended Data Fig. 9a, b). AST remained within normal ranges in both NANITE- and Cas9 RNP-plasmid mice (Extended Data Fig. 9c), and ALT returned to control levels by day 14 (Extended Data Fig. 9d). No significant differences were observed in other biochemical markers (Extended Data Fig. 9e – m). NANITE-plasmids elicited similar neutralizing antibodies as those detected after direct miniEDV administration (3.2 × 10^10^ or 5.5 × 10^10^ miniEDVs per mouse) (Supp. Fig. 9a, b). This humoral immunity did not produce detectable hepatotoxicity (Extended Data Fig. 8b-n). Taken together, these results show that NANITE amplified editing *in vivo* without toxicity.

Lastly, we tested whether NANITE-mediated genome editing amplification *in vivo* requires intercellular spread. Our cell culture data showed that NANITE-mediated amplification depends on endocytosis and endosomal acidification, consistent with VSVG-mediated cell entry of the secreted vesicles (Extended Data Fig. 3e). Previous studies showed that a P127D mutation in VSVG prevents it from acting as a fusogen, abolishing viral cell entry and infection^54^. We introduced this mutation into the VSVG portion of NANITE and confirmed that non-fusogenic NANITE was unable to amplify genome editing in cell culture (Fig. 5a). Non-fusogenic NANITE (VSVG P127D) plasmids behaved identically to the Cas9 RNP-plasmid controls, despite producing vesicles at similar titers to those measured for wildtype NANITE (Fig. 5b). Western blotting of vesicles showed that NANITE (VSVG P127D) produced vesicles harboring both Cas9 and VSVG (Fig. 5c). These results show that *in situ* production of non-fusogenic Cas9-carrying vesicles abolished amplification of genome editing.

**Figure 5.**
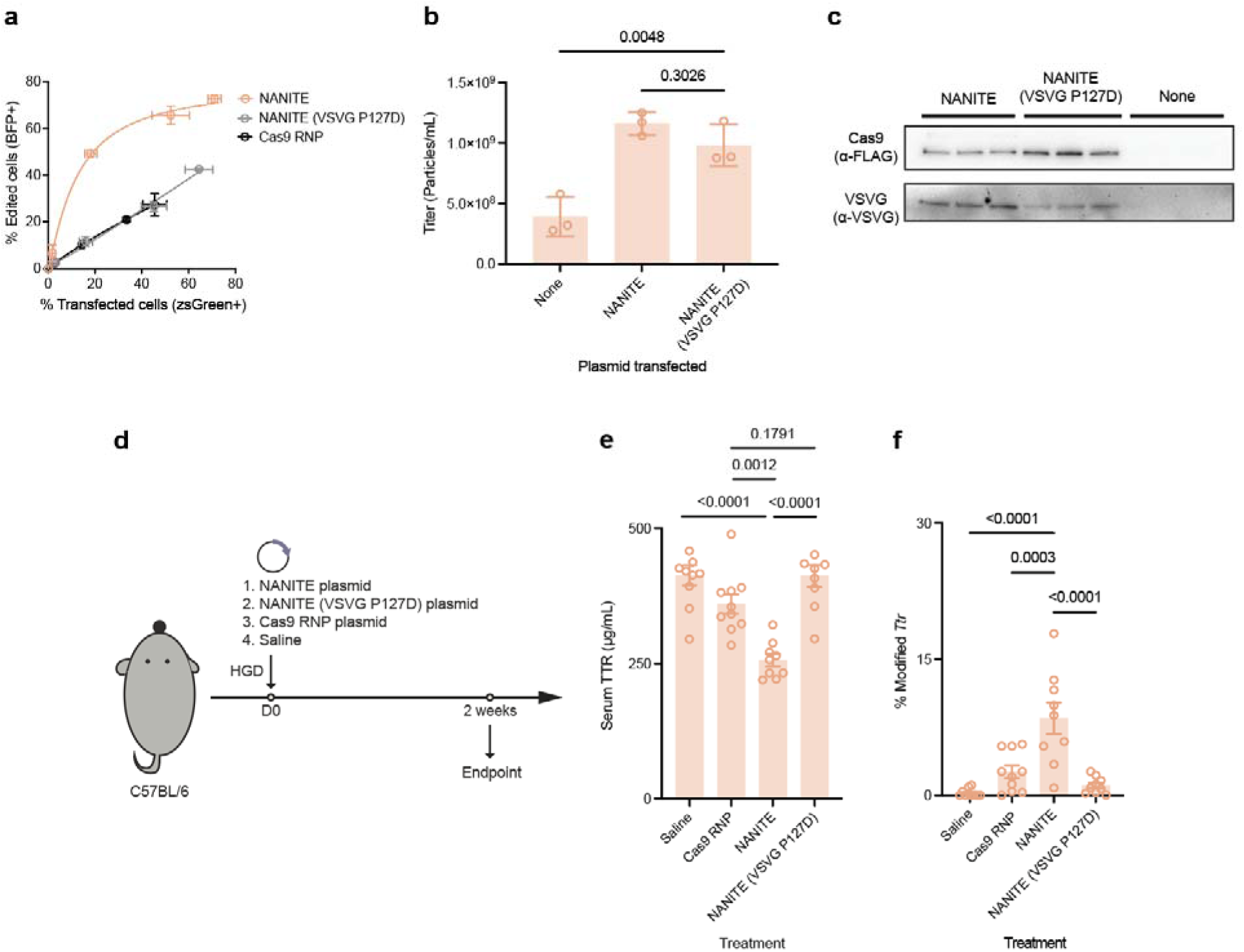
*In vivo* amplification of genome editing requires intercellular spread. **(a)** Fraction of edited cells relative to transfected cells following transfection with NANITE, NANITE (VSVG P127D) or Cas9 RNP plasmids. A P127D mutation in the VSVG portion of NANITE renders VSVG defective and prevents NANITE from spreading and increasing editing. Data are presented as mean ± SD of three replicates. **(b)** Vesicle titers 48 hours after transfection measured by nanoparticle flow cytometry. NANITE and NANITE (VSVG P127D) vesicles were produced similarly. Statistical significance was determined by one-way ANOVA with Dunnett’ multiple comparisons tests. p-values are as indicated. Data are presented as mean ± SD of three biological replicates. **(c)** Western blot of vesicles. Both NANITE and NANITE (VSVG P127D) vesicles carried Cas9 and VSVG proteins. Each lane represents an independent biological replicate. **(d)** Experimental scheme for mouse experiments comparing NANITE with non-fusogenic NANITE (VSVG P127D). Mice were hydrodynamically injected (HGD) with plasmids. Editing was quantified by TTR ELISA on blood serum and sequencing liver genomic DNA two weeks post-injection. **(e)** Serum TTR concentrations. NANITE significantly reduced TTR levels compared to saline, Cas9 RNP and NANITE (VSVG P127D) control groups. Non-fusogenic NANITE (VSVG P127D) was not significantly different compared to Cas9 RNP. **(f)** Sequencing of liver genomic DNA. NANITE-plasmids had more editing than NANITE (VSVG P127D)-, or Cas9 RNP-plasmid groups. Statistical significance in e and f were determined by one-way ANOVA with Tukey’s multiple comparisons test. Relevant p-values are as listed. Data are presented as mean ± SEM (n = 10 for saline and Cas9 RNP, n = 9 for NANITE and NANITE VSVG P127D).

We then compared NANITE-, NANITE (VSVG P127D)- and Cas9 RNP-plasmids in mice. We hydrodynamically injected equimolar doses of each plasmid into mice and collected blood serum and liver DNA two weeks after injection (Fig. 5d). Consistent with our hypothesized mechanism, mice injected with the non-fusogenic NANITE plasmids had similar serum TTR levels to the saline or Cas9 RNP plasmid groups, failing to amplify genome editing (Fig. 5e, f). Altogether these results support the mechanism that NANITE-amplified genome editing requires the ability to spread editing enzymes intercellularly.

## Discussion

A major goal in therapeutic genome editing is ensuring that sufficient cells are edited to produce physiological effects. Research has largely focused on engineering delivery strategies to surpass these thresholds by directly delivering to more cells. Two approaches have been described to increase the number of edited cells after delivery, but they do so indirectly by conferring fitness advantages onto edited cells to stimulate their expansion via cell-type-specific clonal expansion^2^ or selective elimination of unedited cells^3,4^. These approaches lack generalizability and raise safety concerns from depleting healthy cells.

Here, we describe a strategy for directly increasing the number of edited cells post-delivery by enabling transient spread of editing enzymes. This NANoparticle Induced Transfer of Enzymes (NANITE) technology packages editing enzymes into non-replicative vesicles produced by initially transfected cells to transfer into neighboring cells. Nucleic acids encoding NANITE provide transfected cells with all components necessary to transiently edit neighboring cells. NANITE functions across multiple cell types (including HEK-293T, HepG2, and C2C12 cells, and mouse liver cells *in vivo*) without additional engineering. *In vivo*, nucleic acids encoding NANITE tripled genome editing levels compared with Cas9 RNP controls and reduced serum TTR levels by ∼50%, hitting the threshold for improvement and stabilization in transthyretin amyloidosis^55^. Notably, NANITE achieved this despite low initial transfection rates. Cas9 RNP plasmids delivered by the same method did not produce statistically significant reductions in serum TTR. The amplified editing was not toxic *in vivo* and not stimulated by loss of healthy cells, as we detected no inflammation or aberrant cell proliferation in mouse liver.

NANITE creates a sender-receiver system in which cells transfected with NANITE-plasmids become senders of editing enzymes to receiver neighboring cells. Previous sender-receiver systems (including SPIT^15^, NTVE^9^, COURIER^7^ and CROSS-FIRE^10^) were not engineered to amplify genome editing *in vitro* and *in vivo*. SPIT is an engineered cell product, where engineered HEK-293T cells act as a primary delivery vehicle to deliver Cre recombinase *in vivo.* NTVE and COURIER are primarily transcriptomic tools with demonstrated *in vitro* transfer of mRNA from HEK-293T sender cells. CROSS-FIRE is an exosome-based system that transfers sgRNAs to recipient cells stably expressing Cas9 *in vitro*.^10^ NANITE creates a system that transfers Cas9 RNPs. The results presented here show that genome editing can be amplified post-delivery *in vitro* and *in vivo* using nucleic acid payloads that encode a sender-receiver system.

Results using a non-fusogenic form of NANITE demonstrated that amplified *in vivo* genome editing requires intercellular spread. We could not quantify transfection and editing simultaneously *in vivo* because transfection declined within 24 hours while editing took longer^56^. Although NANITE is currently formulated as a plasmid because it requires a cassette expressing sgRNA to form Cas9 RNPs for vesicle encapsulation, future advances including conversion to an all-RNA format could expand the delivery vehicles and target tissues available for *in vivo* editing^57–60^. It may also be possible to formulate NANITE with alternative genome editors including base- and prime-editing versions of Cas9, though differences in exposure time between transfected and receiver cells will need to be considered. In the future, we envision pairing tissue-tropic delivery vehicles with correspondingly targeted NANITE constructs to achieve tissue-specific editing amplification. By increasing the number of cells that can be edited post-delivery, NANITE increases the likelihood of achieving effective *in vivo* genome editing for a wide range of therapeutic applications.

## Extended Data Figures

**Extended Data Figure 1.**
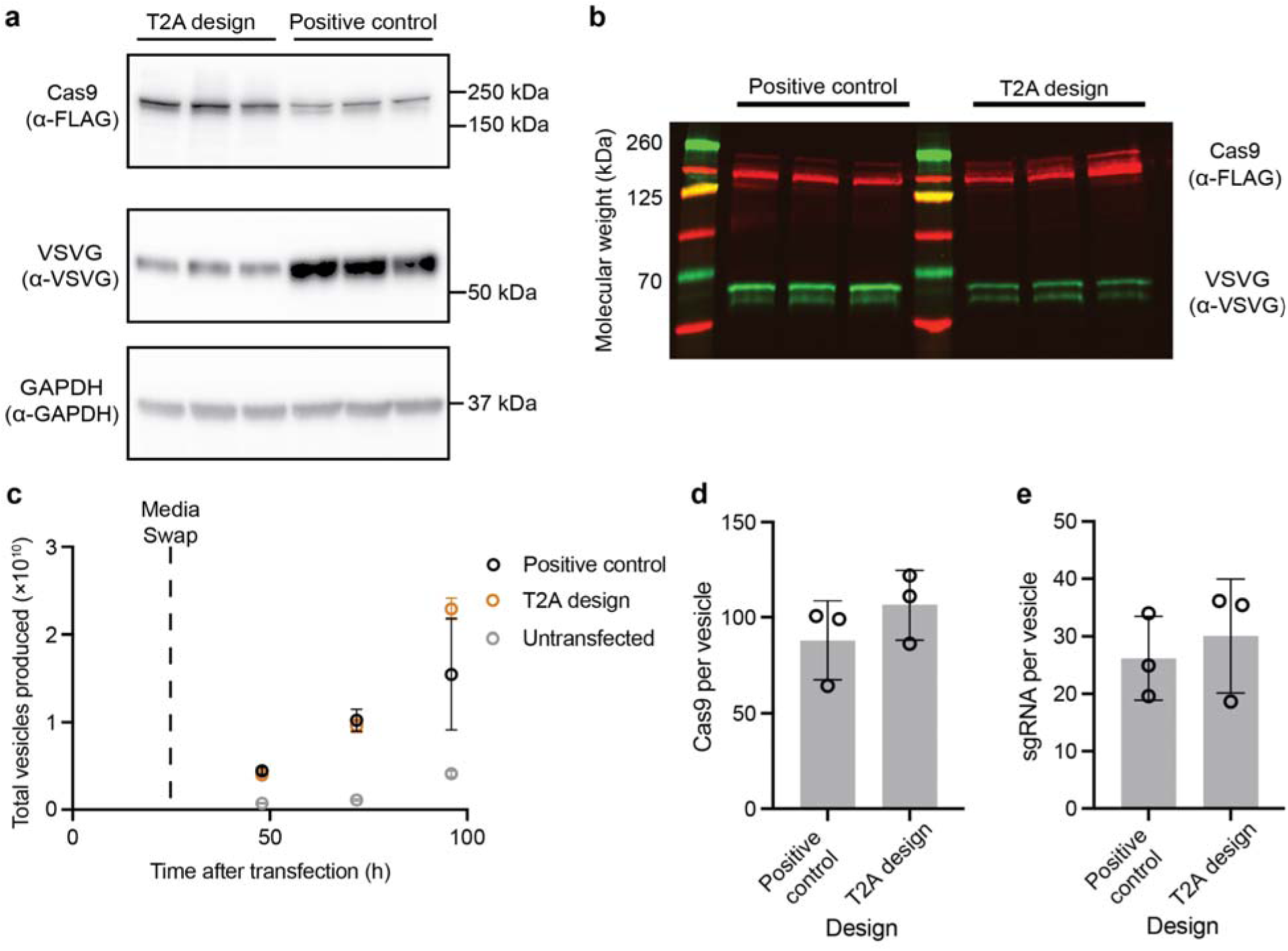
Characterization of T2A design. **(a)** Western blot of cell lysates 24 hours post-transfection with T2A or positive control plasmids. The T2A design resulted in complete cleavage of VSVG. The miniGag-SpCas9 fusion protein (C-terminally FLAG-tagged) migrates at approximately 210 kDa as expected. VSVG migrates at approximately 55 kDa a expected. GAPDH serves as a loading control. Each lane represents an independent biological replicate. **(b)** Western blot of vesicles. Less VSVG is present on NANITE vesicles compared to the miniEDVs. Each lane represents an independent biological replicate. **(c)** Rate of vesicle production after transfection with positive control or T2A design plasmids. Vesicles were produced at a similar rate as measured by nanoparticle flow cytometry. Data are presented as mean ± SD of three biological replicates. **(d)** Amount of Cas9 packaged per vesicle as quantified by ELISA. The T2A design packaged a similar amount of Cas9 per vesicle as compared to the positive control. Data are presented as mean ± SD of three biological replicates. **(e)** Amount of sgRNA packaged per vesicle as quantified by qRT-PCR. The T2A design packaged a similar amount of sgRNA per vesicle as the positive control. Data are presented as mean ± SD of three biological replicates.

**Extended Data Figure 2.**
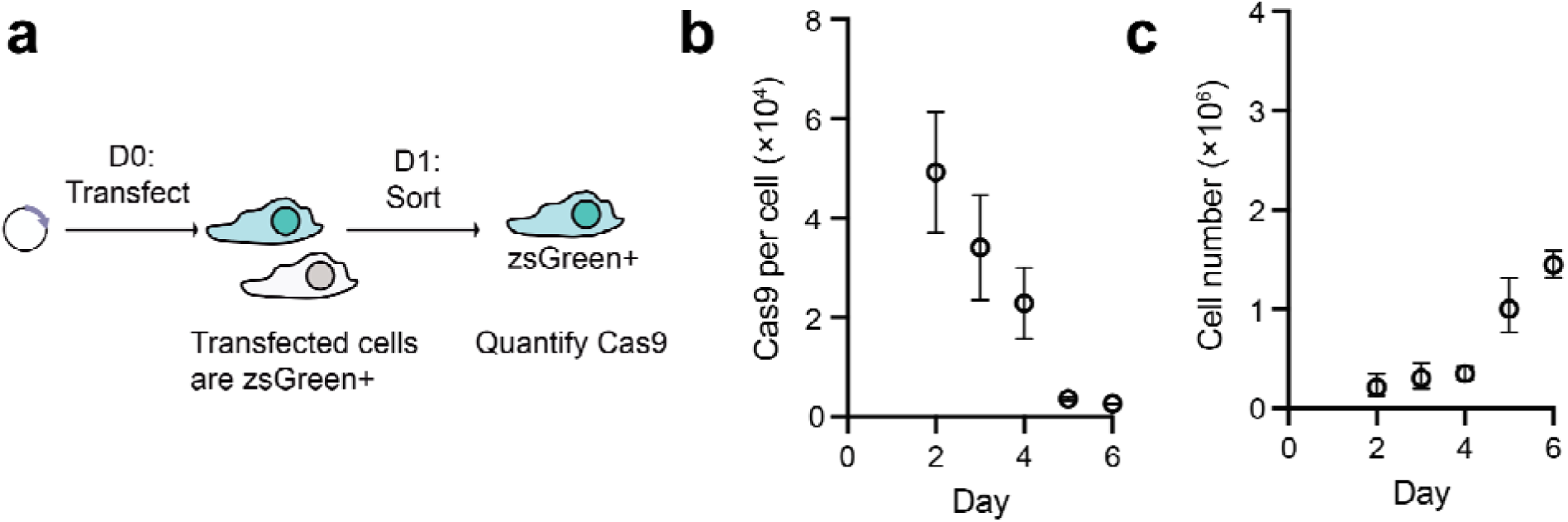
NANITE decreases Cas9 concentrations in transfected cells over time. **(a)** We isolated cells one day after transfection using fluorescence activated cell sorting, then quantified intracellular Cas9 concentrations using ELISA over time. **(b)** Cas9 per cell as determined using ELISA. Cas9 levels declined gradually. **(c)** The number of cells over time. Data are presented as mean ± SD of three biological replicates.

**Extended Data Figure 3.**
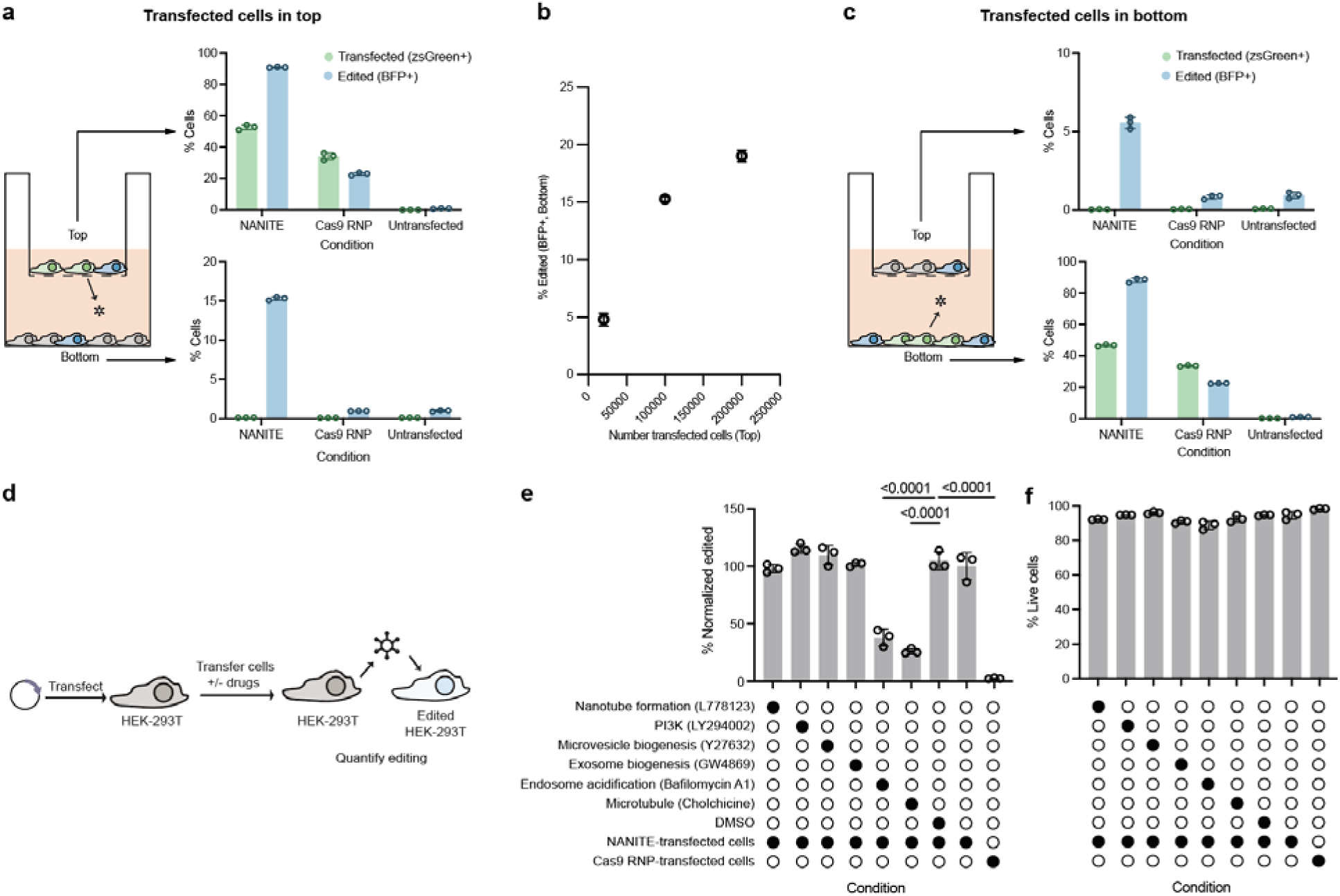
NANITE spreads by vesicles. **(a)** Transfected cells were seeded in the top well. Edited cells in the top and bottom wells were quantified by flow cytometry. Edited cells were detected in the bottom well after transfection with NANITE-plasmids. **(b)** Fraction of edited cells in the bottom well as a function of the number of transfected cells seeded in the top well. **(c)** Transfected cells were seeded in the bottom well. Edited cells in the top and bottom wells were quantified by flow cytometry. Edited cells were detected in the top well after transfection with NANITE-plasmids. **(d)** Schematic of the inhibitor experiment. HEK-293T cells were transfected with either NANITE- or Cas9 RNP-plasmids and transferred 24 h after transfection into receiver cells with inhibitors. **(e)** Edited cells quantified by flow cytometry. The inhibited pathway is indicated with the drug. Only inhibitors of different steps in the endocytic pathway (bafilomycin A1 and colchicine) significantly reduced transfer and editing. Editing values were normalized to the NANITE-plasmid condition with no inhibitors. Statistical significance was determined using ordinary one-way ANOVA with Dunnett’s multiple-comparison test against the untreated NANITE condition. Significant p-values are indicated. **(f)** Cell viability as determined by flow cytometry after staining with Zombie NIR live/dead stain. The inhibitor concentrations used did not affect cell viability. Schematics are not to scale. All data are presented as mean ± SD from three biological replicates.

**Extended Data Figure 4.**
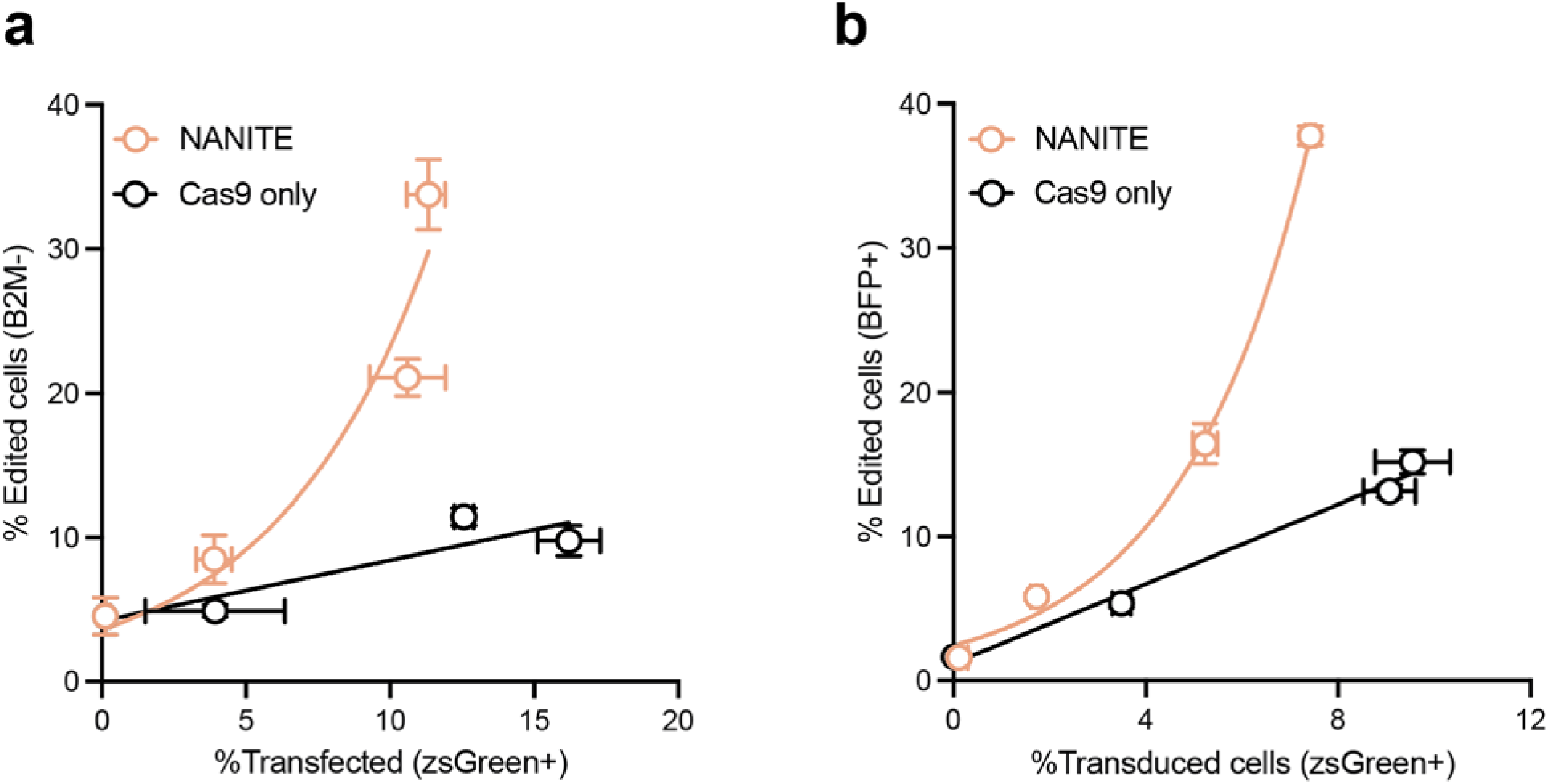
NANITE functions in other cell types. **(a)** Percentage of edited HepG2 cells at the endogenous *B2M* locus relative to transfected cells. **(b)** Percentage of edited BFP-reporter C2C12 cells relative to transfected cells. Data are presented as mean ± SD of three biological replicates.

**Extended Data Figure 5.**
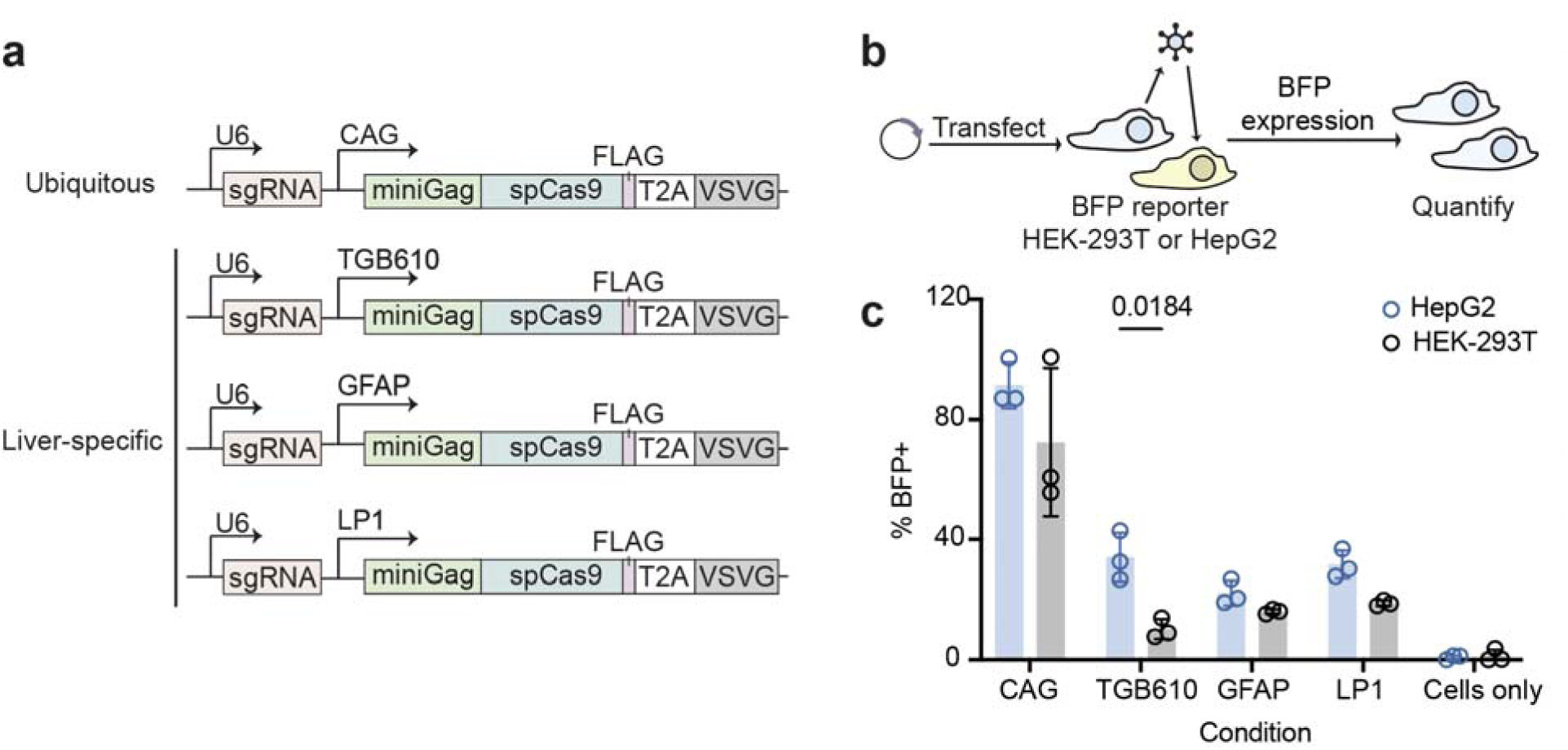
NANITE tropism can be controlled with cell-specific promoters. **(a)** Schematics of NANITE plasmids with liver-specific promoters, where the CAG promoter i replaced with either the TBG610, GFAP, or LP1 promoter. **(b)** Experimental scheme to test promoters. BFP-reporter HEK-293T or HepG2 cells were transfected with plasmids. NANITE expression and activity result in BFP expression, which was quantified. Schematics are not to scale. **(c)** Percentage of BFP+ HEK-293T or HepG2 cells. The TBG promoter showed significantly higher NANITE activity in HepG2 cells compared to HEK-293T cells. Statistical significance was determined by two-way ANOVA with Šidák’s multiple comparisons test. Significant p-values are indicated. Data are presented as mean ± SD of three biological replicates.

**Extended Data Figure 6.**
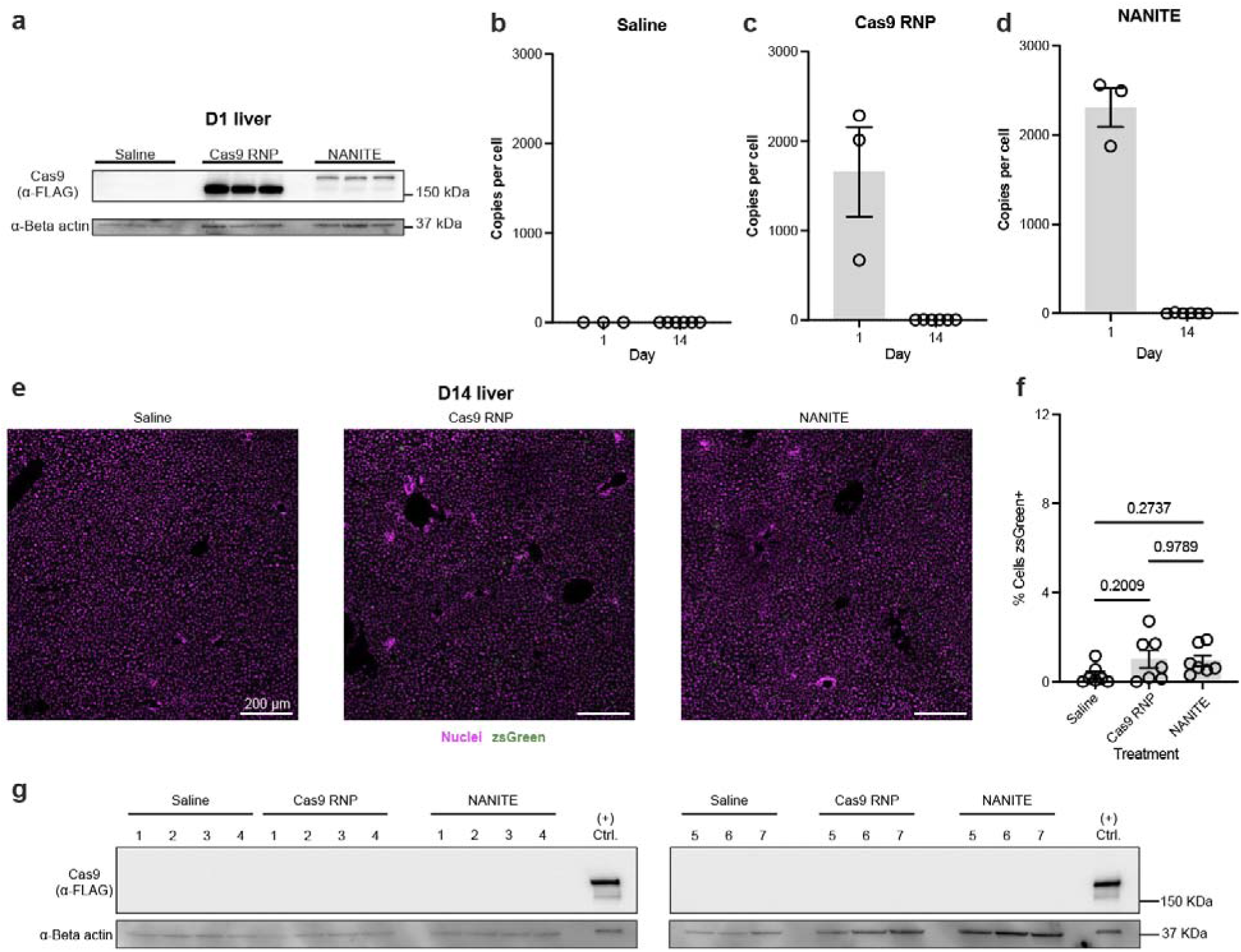
*In vivo* persistence of NANITE- and Cas9 RNP-plasmids. **(a)** Western blot of Cas9 and NANITE proteins in liver lysates one day post-injection, detected with anti-FLAG antibodies. β-actin serves as a loading control. Each lane represents a technical replicate. Experiments were repeated twice with similar results. **(b)** Copies of plasmids quantified by qPCR in saline-treated mice livers. Data are presented as the mean ± SEM of three mice on day 1 and seven mice on day 14. **(c)** Copies of plasmids quantified by qPCR in Cas9 RNP plasmid-treated mice livers. Data are presented as the mean ± SEM of three mice on day 1 and seven mice on day 14. **(d)** Copies of plasmids quantified by qPCR in NANITE plasmid-treated mice livers. Data are presented as the mean ± SEM of three mice on day 1 and seven mice on day 14. **(e)** Representative immunofluorescence images of liver sections harvested 14 days post-injection. Nuclei are stained with DAPI (magenta). zsGreen is shown in green. No zsGreen expression is detected on day 14. Scale bar as indicated. **(f)** Percentage of zsGreen+ liver cells quantified using CellProfiler from images in (e). One field of view per mouse was analyzed. Data are presented as the mean ± SEM of seven mice. Statistical significance was determined by one-way ANOVA with Tukey’s multiple comparison test. No significant differences were observed between saline, Cas9 RNP-plasmid, or NANITE-plasmid groups. **(g)** Western blot of Cas9 and NANITE proteins in liver lysates, detected with anti-FLAG antibodies. No Cas9 RNP or NANITE expression was detected at day 14. β-actin serves as a loading control. Each lane represents lysate from one mouse numbered one to seven.

**Extended Data Figure 7.**
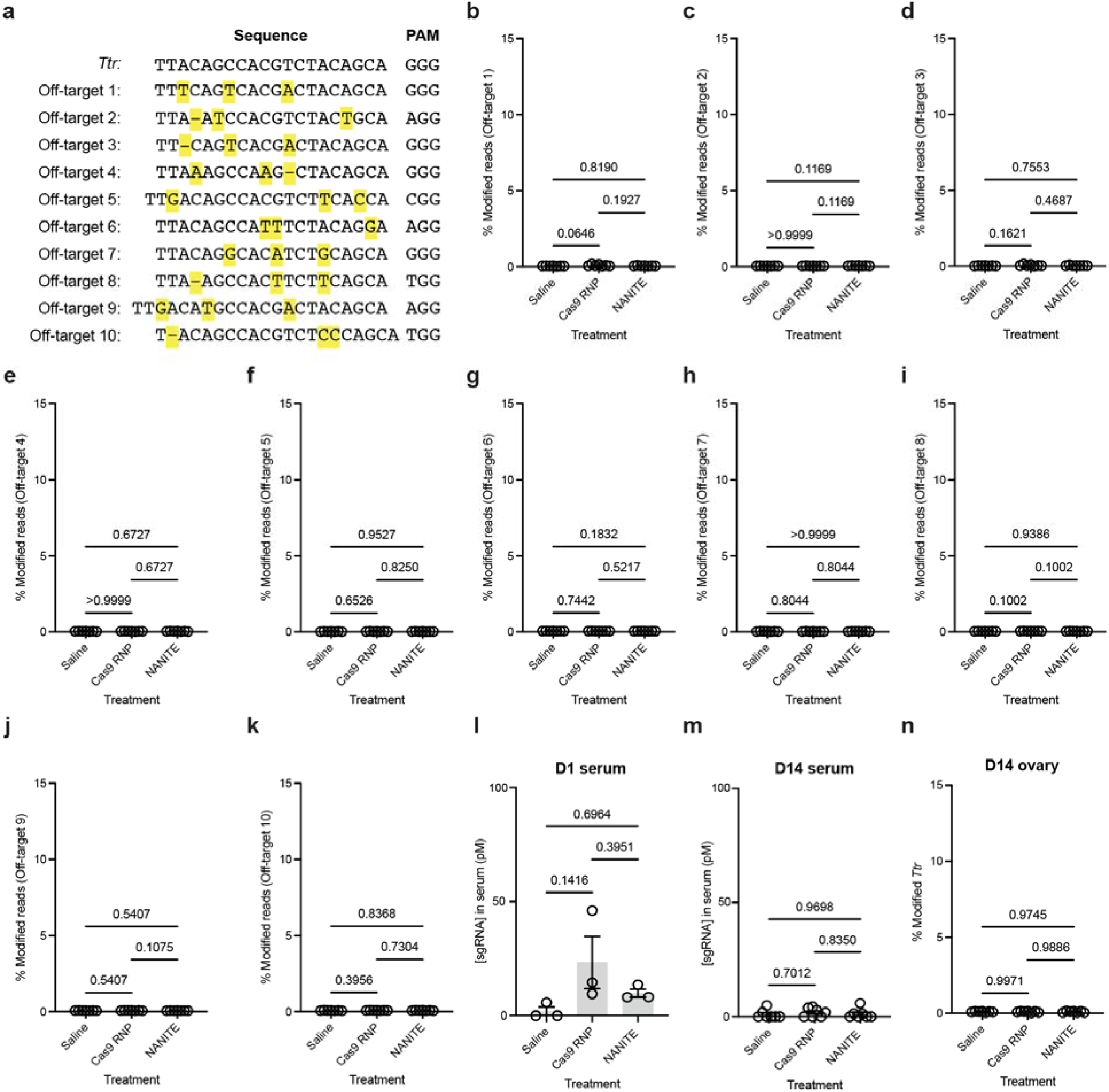
NANITE did not elevate off-target editing. **(a)** Off-target loci predicted by Cas-OFFinder. Mismatches relative to the on-target sequence are highlighted. Th protospacer adjacent motif (PAM) is indicated. **(b-k)** Percentage of modified reads at the indicated off-target sites in liver genomic DNA harvested on day 14. **(l)** Serum sgRNA concentration on day 1. Data are presented as the mean ± SEM of three mice. **(m)** Serum sgRNA concentration on day 14. For (l) and (m), samples below the limit of detection are plotted on the x-axis. **(n)** Percentage of modified reads at the *Ttr* locus in ovary genomic DNA harvested on day 14. Data are presented as the mean ± SEM of seven mice unless otherwise indicated. Statistical significance was determined by one-way ANOVA with Tukey’s multiple comparison test. No significant differences were observed between treatment groups.

**Extended Data Figure 8.**
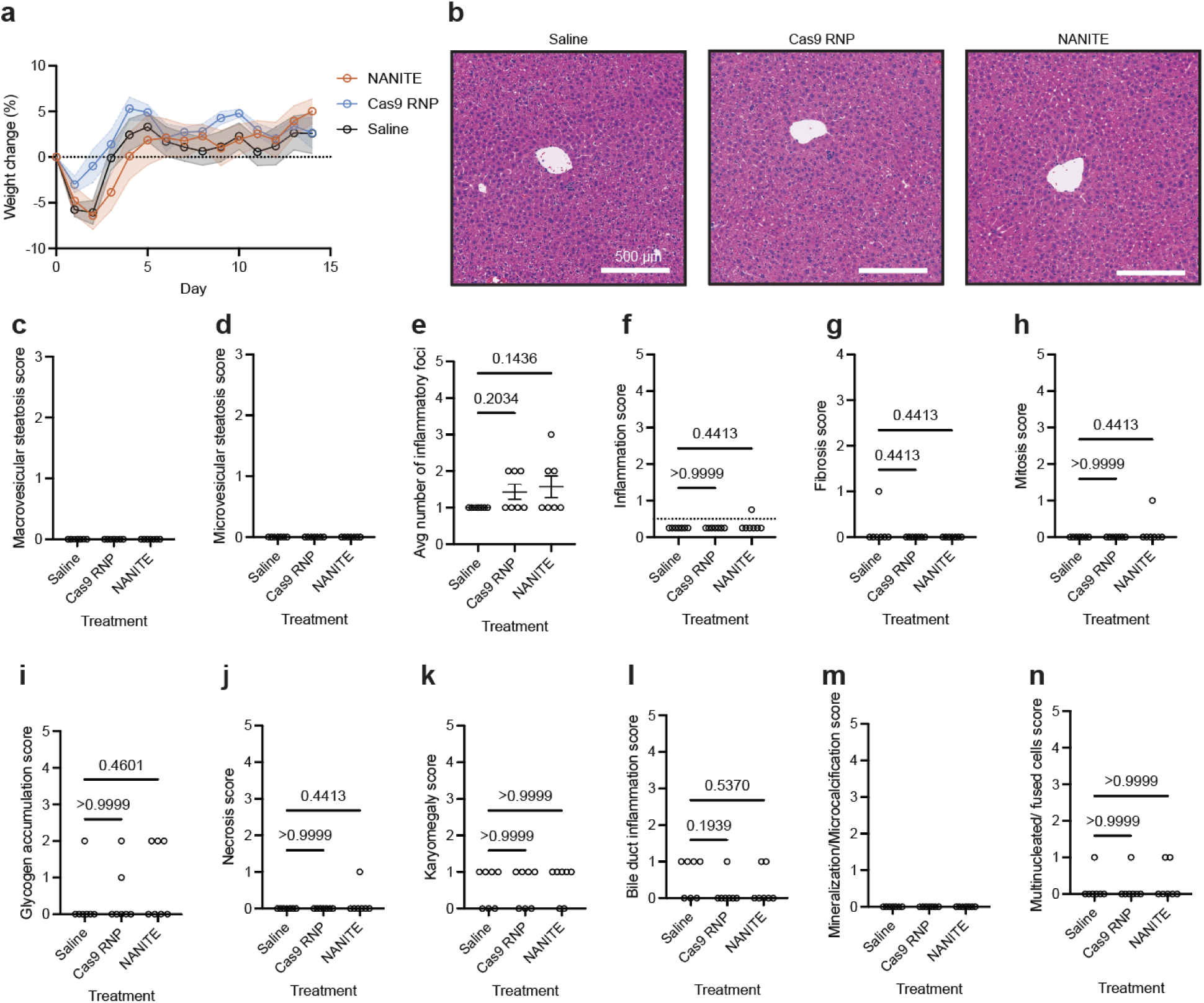
NANITE did not adversely affect mouse health. **(a)** Body weight changes following treatment. Data are presented as the mean ± SEM of seven mice. Statistical significance was determined by two-way ANOVA with Tukey’s multiple comparison test. No significant differences were observed between groups at any time point. **(b)** Representative hematoxylin and eosin-stained liver sections. A subsection from a larger slide is shown. Scale bar as indicated. **(c)** Macrovesicular steatosis score assessed on whole liver slides. A score of zero indicates <5% tissue affected. **(d)** Microvesicular steatosis score. A score of zero indicates <5% tissue affected. **(e)** Average number of inflammatory foci per slide. Data are presented a the mean ± SEM of seven mice. **(f)** Inflammation score. A score of <0.5 indicates normal physiology (dotted line). A score of 0.5 - 1 indicates slight inflammation **(g)** Fibrosis score. Zero indicates no fibrosis. Four indicates cirrhosis. **(h)** Mitosis score. **(i)** Glycogen accumulation score. **(j)** Necrosis score. **(k)** Karyomegaly score. **(l)** Bile duct inflammation score. **(m)** Mineralization score. (**n)** Multinucleated/fused cell score. For (h–n), a score of 0 indicates no tissue affected. A score of 1 indicates 1–10% affected. A score of 2 indicates 11-25% tissue affected. All scores (c–n) were assessed by a certified pathologist in a blinded manner. Statistical significance was determined using a non-parametric Kruskal-Wallis test with Dunn’s multiple comparison test to saline treated animals. Each data point represents one mouse.

**Extended Data Figure 9.**
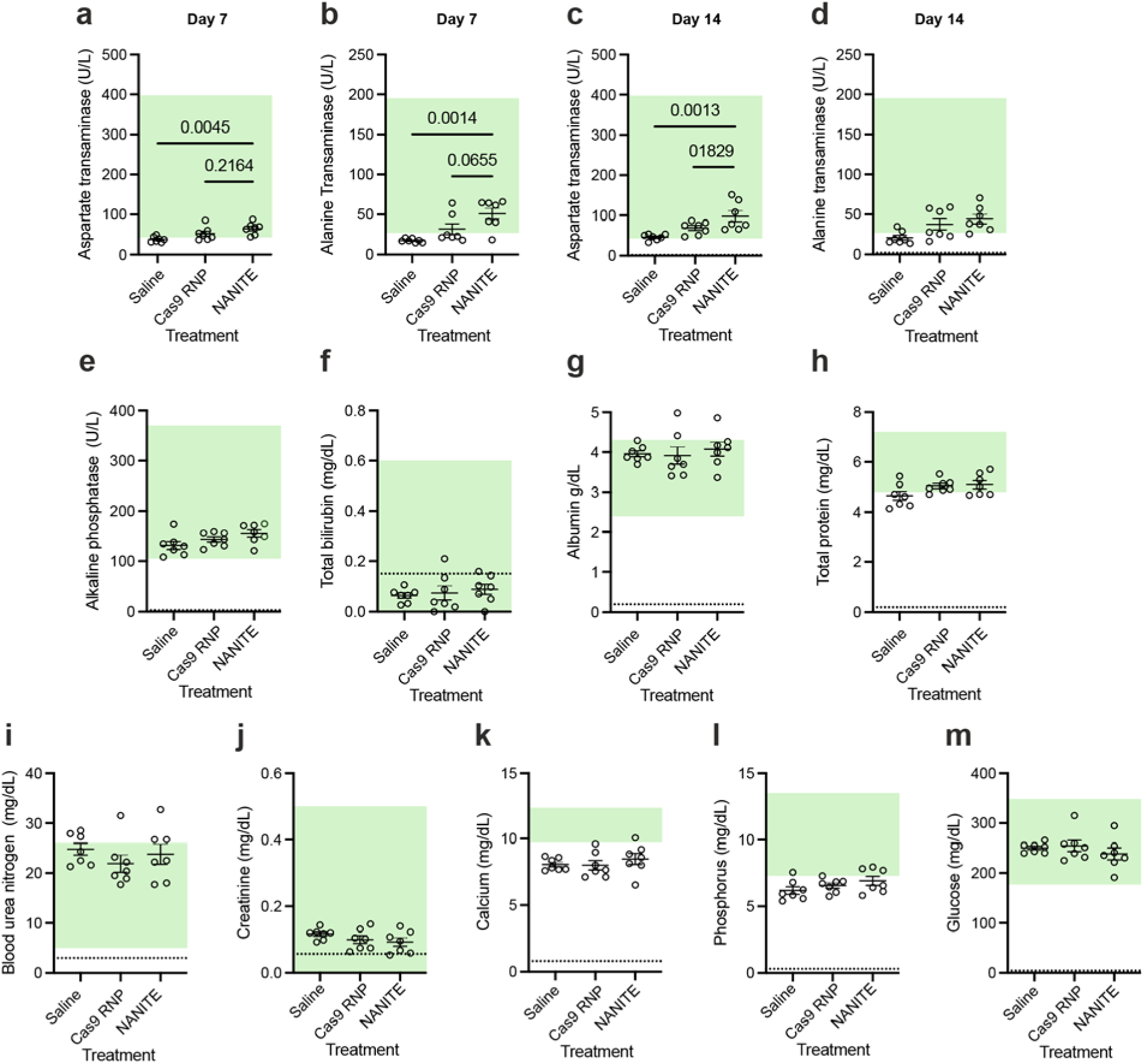
Blood Biochemistry of mice treated with NANITE-plasmids. Serum concentrations of **(a)** Aspartate transaminase on day 7, **(b)** Alanine transaminase on day 7, **(c)** Aspartate transaminase on day 14, **(d)** Alanine transaminase on day 14. **(e)** Alkaline phosphatase, **(f)** Total bilirubin, **(g)** Albumin, **(h)** Total protein, **(i)** Blood urea nitrogen, **(j)** Creatinine, **(k)** Calcium, **(l)** Phosphorus, and **(m)** Glucose. In all panels, the green shaded area indicates the normal physiological range. Dotted lines indicate detection limits where appropriate. Data are presented as the mean ± SEM of seven mice. Each point represents one mouse. Statistical significance was determined by one-way ANOVA with Tukey’s multiple comparison test.

## Methods

### Cloning plasmids

Plasmids were cloned using HiFi assembly (New England Biolabs, #E2621X). DNA fragments for assembly were generated by PCR using Q5 PCR Master Mix (New England Biolabs, #M0492L) or ordered as gene blocks (Twist Bioscience or Integrated DNA Technologies). To create the NANITE plasmid, VSVG was amplified from pMD2.G (Addgene, #12259) using primers with appropriate overhangs (Integrated DNA Technologies). The VSVG and T2A fragments (Integrated DNA Technologies) were inserted downstream of Cas9 in pWN_U6-B2M-miniGag (Addgene, #228957). Other plasmids were cloned using a similar strategy. CRISPR-Cas9 spacer sequences are listed in Table S1.

Plasmid and HiFi assemblies were transformed into Mach1 *E. coli* cells (Thermo Fisher Scientific, #C862003) rendered competent using the Mix & Go *E. coli* Transformation Kit (Zymo Research, #T3001). Mach1 cells were grown with appropriate antibiotic selection, then plasmids were extracted and purified using miniprep (Qiagen, #27104), midiprep (Zymo Research, #D4200), or maxiprep kits (Zymo Research, #D4202). For animal experiments, plasmids were extracted using the EndoFree Plasmid Maxi Kit (Qiagen, #12362) to minimize endotoxin contamination. Endotoxin levels were quantified using limulus amebocyte lysate assays (Thermo Fisher Scientific, #A39552S) prior to administration. All plasmids were verified by whole-plasmid sequencing (Plasmidsaurus).

### Cell culture

HEK-293T cells (gift from Dr. Melanie Ott, Gladstone Institutes) were seeded on 15-cm plates (Fisher Scientific, #FB012925) in DMEM (Corning, #10-013-CV) supplemented with 10 v/v% FBS (Avantor, #97068-085) and 1 v/v% penicillin/streptomycin (Thermo Fisher Scientific, #15140122). Cells were incubated at 37°C with 5% CO and passaged as required using 0.25% trypsin, with 0.1% EDTA (Corning, #MT25053CI). C2C12 cells (ATCC, #CRL-1772) were similarly cultured. HepG2 cells (ATCC, #HB-8065) were cultured in EMEM (ATCC, #30-2003) supplemented with 10 v/v% FBS and 1 v/v% penicillin /streptomycin. All cells were routinely tested for mycoplasma contamination (Stem Cell Core, Gladstone Institutes).

### Generating BFP-reporter cells

EF-1α promoter, blue fluorescent protein (BFP) reporter, and puromycin resistance fragments (Fig. S1a) were cloned into a pLVX backbone using HiFi assembly to generate a lentiviral transfer plasmid. To produce lentiviral vectors, 4 million HEK-293T cells were seeded on 10-cm dishes (Corning, #35300). The following day, cells were transfected with 2.5 μg of transfer plasmid, 1 μg pMD2.G, and 10 μg psPAX2 using polyethylenimine (PEI; Polysciences, #23966) at a 3:1 ratio (PEI:DNA by mass). Excess plasmid was removed after 24 hours, and lentiviral vectors were harvested at 48 hours. HEK-293T cells (48,000 cells) were incubated with 0.5 mL lentiviral vectors and 8 μg/mL polybrene (MilliporeSigma, #H9268) for 48 hours, followed by selection in 2 μg/mL puromycin (InvivoGen, #ant-pr-1) for 48 hours.

To validate reporter cells, miniEDVs targeting either the BFP-reporter or *TRAC* locus were produced as previously published^18^, then 8–500 μL were incubated with 15,000 reporter cells in 24-well plates (Corning, #353047). Cells were trypsinized after 72 hours and stained with 1 μL Zombie NIR live-dead stain (BioLegend, #423106). BFP expression was quantified by flow cytometry on an Attune NxT Flow Cytometer (Thermo Fisher Scientific). Flow gating is shown in Fig. S1d,e. Genomic DNA was harvested using QuickExtract reagent (Biosearch Technologies, #QE09050). The reporter locus was amplified by PCR (BFPreporter.rev and BFPreporter.for; sequences in Table S2) and Sanger sequenced (Elim Biopharm Inc.). Insertion and deletion frequencies were calculated using the TIDE analysis webserver^61,62^. Cells were stored in liquid nitrogen for use in downstream experiments. Similar procedures were used to generate BFP-reporter C2C12 and HepG2 cells.

### Testing single plasmids encoding NANITE

To produce vesicles, 600,000 HEK-293T cells were seeded per well of a 6-well plate (Corning, #3516). The following day, cells were transfected with 2.2 μg NANITE plasmid or 2 μg pWN_U6-BFPreporter-miniGag and 0.2 μg pMD2.G as a positive control using PEI. Excess plasmid was removed after 24 hours. Vesicles were harvested 48 hours post-transfection as previously published^18^, and up to 50 μL was incubated with 7,500 BFP-reporter HEK-293T cells in 48-well plates (Corning, #3548). Cells were trypsinized after 72 hours and stained with Zombie NIR live-dead stain (BioLegend, #423106). BFP expression was quantified by flow cytometry on an Attune NxT Flow Cytometer. Flow gating is shown in Fig. S1d, e.

For Western blotting of cell lysates, HEK-293T cells were harvested 48 hours post-transfection using RIPA lysis buffer (Thermo Fisher Scientific, #89900) with 1× protease inhibitors (Thermo Fisher Scientific, #78429). Lysates were centrifuged at >19,000 × g for 5 minutes to remove insoluble aggregates. Supernatants were stored at −80°C. Protein concentrations were determined using the BCA assay (Thermo Fisher Scientific, #23225). Samples normalized for protein concentration were mixed with 4× Laemmli sample buffer (Bio-Rad, #1610747) to a final 1× concentration with 1% (v/v) 2-mercaptoethanol (MilliporeSigma, #M3148), then denatured at 90°C for 5 minutes. Samples were run on SDS-PAGE gels (Bio-Rad, #5678094) for 2 hours at 120 V at 4°C. Proteins were transferred to 0.2 μm nitrocellulose membranes using a Trans-Blot Turbo Transfer Pack (Bio-Rad, #1704159) at 2.5 A and 25 V for 7 minutes using a Trans-Blot Turbo Transfer System (Bio-Rad). Membranes were blocked with 5 w/v% non-fat dry milk (Lab Scientific bioKEMIX, #M0841) in 1× TBS-T for at least 2 hours at room temperature. 1×TBS-T consists of 10 mM Tris (Corning, #46-030-CM), 150 mM NaCl (Thermo Scientific, #J21618-A9), and 0.05 v/v% Tween-20 (Fisher, #BP337). Membranes were incubated overnight at 4°C with anti-FLAG antibody (MilliporeSigma, #F1804, 1:1,000), anti-VSVG antibody (Kerafast, #Kf-Ab01401-23.0, 1:1,000), or anti-GAPDH antibody (Santa Cruz Biotechnology, #sc-365062, 1:1,000) in blocking solution as appropriate. Membranes were washed 3–5 times for 5 minutes each with 1× TBS-T, then incubated with the appropriate secondary antibody (goat anti-mouse, Invitrogen, #62-6520, 1:1,000; or goat anti-rabbit, Invitrogen, #656120, 1:1,000) in blocking solution for 1 hour at room temperature. Membranes were washed 3–5 times and developed using chemiluminescent substrate (Thermo Fisher Scientific, #34580). Images were acquired using a ChemiDoc MP imaging system (Bio-Rad). The unprocessed blots are shown in Fig. S10.

### Characterizing NANITE

Protease protection assays were performed as published^8^. Vesicles were produced in HEK-293T cells and harvested after ∼72 hours as described above. Samples were incubated on ice for 30 minutes alone or with 1 v/v% Triton X-100 (Sigma-Aldrich, #X100), 10 μg/mL proteinase K (New England Biolabs, #P8107S), or both. PMSF (Sigma-Aldrich, #P7626) was added to a final concentration of 5 mM to inhibit the proteinase K and incubated for 5 minutes on ice. Laemmli buffer was added to a final concentration of 20 v/v% and samples were heated at 98°C for 5 minutes. Samples were run on SDS-PAGE gels (Bio-Rad, #4561095) and transferred to PVDF membranes (Bio-Rad, #1620177) at 90 V and 4°C for 1 hour in transfer buffer. The transfer buffer consists of 25 mM Tris (Avantor, #4099-06), 192 mM glycine (Bio-Rad, #1610724), and 20 v/v% methanol (VWR, #BDH1135). Membranes were blocked for 1 hour at room temperature on an orbital shaker in blocking buffer consisting of 1× PBS (Gibco, #14190) with 0.1% Tween-20 (Sigma-Aldrich, #P7949) saturated with non-fat dry milk (Bio-Rad, #1706404XTU). Membranes were washed four times with 1྾PBS containing 0.1% Tween-20 (PBS-T), then incubated overnight at 4°C in blocking buffer with anti-FLAG antibody (Sigma-Aldrich, #F1804, 1:2,000). Membranes were washed four times with PBS-T, then incubated with secondary antibody (LI-COR, #926-68070, 1:20,000) for 1 hour at room temperature. The membranes were then washed four times with PBS-T and imaged on a LICOR imager (LICORBio). The unprocessed blot is shown in Fig. S10.

To compare Cas9 and VSV-G levels in vesicles produced from cells transfected with NANITE versus the two-plasmid control, vesicles were produced as described above and analyzed qualitatively by Western blot. 30 µL of each sample was run on a Bolt 4-12% Bis-Tris Plus WedgeWell Gel (invitrogen, #NW04120BOX) and proteins were transferred using an iBlot 3 dry transfer system (Thermo Fisher Scientific, #IB31001) at 25 V for 6 minutes. Antibody staining was performed as described above with the addition of rabbit anti-VSVG (Absolute Antibody, #Ab01401-23.0, 1:1,000) and anti-rabbit secondary antibody (Licor, #926-32211, 1:2,000). The unprocessed blot is shown in Fig. S10.

The concentration and RNP loading of NANITE and two-plasmid control vesicles were quantified as described previously.^18^ Vesicles are produced as described above and 100 µL was harvested at 48, 72, and 96 hours after transfection. To quantify the vesicle numbers, 10 µL of each sample was diluted into at least 140 µL of nanoFCM EV diluent (nanoFCM, #EP-23031), then immediately analyzed using a nanoparticle flow cytometer (nanoFCM, Flow NanoAnalyzer). For Cas9 quantification, 20 µL of vesicles collected at 48 hours was mixed with 180 µL of RIPA buffer supplemented with protease and phosphatase inhibitors. Samples were stored at −80 °C until analysis. The quantity of Cas9 was determined using a Cas9 ELISA Kit (Cell Biolabs, #PRB-5079) following the manufacturers protocol. For sgRNA quantification, 50 µL of supernatant was combined with 50 µL of DNA/RNA Shield Direct stored at −80 °C until analysis. Standard curves of sgRNA were generated using unmodified sgRNA (Integrated DNA Technologies). 4 µL of samples or standards were combined with 2.5 μL of Luna Universal qPCR Master Mix (New England Biolabs, #M3003), 2 μL of reverse-transcription primer, 0.5 μL of probes (Thermo Fisher Scientific, Custom TaqMan Small RNA Assay #CTAACJP), and 1 μL of nuclease-free water (Thermo Fisher Scientific, #AM9937). Reactions were assembled in 384-well plates (Applied Biosystems, #A36931) and sealed with optical adhesive film (Applied Biosystems, #4311971). RT-qPCR was run on a QuantStudio 5 Real-Time PCR System (Applied Biosystems) with the following settings: carryover prevention (25°C, 30 seconds), reverse transcription (55°C, 15 minutes), initial denaturation (95°C, 1 minute), and 45 cycles of denaturation (95°C, 10 seconds) and extension (60°C, 60 seconds).

### In situ editing with NANITE

BFP-reporter HEK-293T cells (10,000 cells/well) were seeded in 96-well plates (Corning, #3596). The following day, cells were transfected with 15–100 ng NANITE or Cas9 RNP control plasmids using PEI at a 3:1 ratio (PEI: DNA by mass). The media were replaced after 24 hours to remove excess plasmids. Flow cytometry was performed at 24, 36, 48, 72, and 96 hours post-transfection as described above, with example gating shown in Fig. S1d and Fig. 2b. Experiments in HepG2 and C2C12 cells were performed similarly using jetPRIME (Sartorius, #101000015) and Cell Avalanche (EZ Biosystems, #EZT-C2C1-1) transfection reagents, respectively. To quantify B2M expression in HepG2 cells, cells were stained with 5 μL APC anti-human β2-microglobulin (BioLegend, #316312) and 1 μL Zombie UV live-dead stain (BioLegend, #423108). B2M expression was quantified on an Attune NxT Flow Cytometer.

To visualize NANITE spread, 20,000 BFP-reporter cells per well were seeded into polylysine-coated (Thermo Fisher Scientific, #A3890401) chamber slides (Thermo Fisher Scientific, #154526). Cells were transfected with NANITE or Cas9 RNP plasmids. Media were replaced after 24 hours to remove excess plasmid. At each timepoint, cells were fixed with 4% buffered formaldehyde (MilliporeSigma, #1.00496), then permeabilized with 0.1 v/v% Triton X-100 (Fisher Scientific, #BP151-100) in 1× PBS. Fixed permeabilized cells were washed with 5 w/v% bovine serum albumin (MilliporeSigma, #A9647) in 1× PBS. Nuclei were stained with RedDot 2 far-red nuclear stain (Biotium, #40061) for 10 minutes at room temperature. Stained cells were washed three times with 5 w/v% bovine serum albumin in 1× PBS before mounting with antifade mounting medium (Vector Laboratories, #VECTH170010). Cells were imaged with identical settings using a Zeiss LSM880 confocal microscope (Gladstone Histology and Light Microscopy Core). Two biological replicates per condition were imaged, with at least three fields of view each. Nuclei were segmented and classified as zsGreen- or BFP-positive using CellProfiler 4.2.8^23^. The distance from each BFP nucleus to the nearest zsGreen nucleus was measured and binned into histograms. Frequency distributions were fitted to a one-phase exponential decay in GraphPad Prism v.10.6.1 to determine half-distances.

### Cas9 concentrations in transfected cells over time

HEK-293T cells were transfected with NANITE plasmids as described above in triplicate. Transfected cells (zsGreen-positive) were isolated 24 hours post-transfection using fluorescence activated cell sorting (BD Biosciences, BD FACS Aria Fusion 2). Following sorting, cells from each replicate were divided equally and seeded into 6-well plates. On each subsequent day (up to day 6), one well of cells were harvested, then counted using a Countess II Automated Cell Counter (Life Technologies). Harvested cells were subsequently washed by centrifugation at 300 × g for 5 min and lysed in RIPA buffer supplemented with protease and phosphatase inhibitor cocktail. Cas9 protein concentrations were quantified using a Cas9 ELISA Kit according to the manufacturer’s instructions.

### Transwell experiments

BFP-reporter HEK-293T cells (1,000,000 cells/well) were seeded in a 6-well plate in 2 mL of complete DMEM. The following day, cells were transfected with either 2,500 ng of NANITE or Cas9 RNP plasmids using PEI as described above. After 24 hours, transfected cells were washed, trypsinized, then 20,000, 100,000, or 200,000 transfected cells were seeded into the top of the Transwell (100 µL) or the bottom (600 µL) of the Transwell system (Corning, #3398) as indicated. The same number of untransfected BFP-reporter cells were seeded in the opposite compartment. Flow cytometry was performed as described above to quantify editing 48 hours after co-culture.

### Small molecule inhibitor experiments

Colchicine (Sigma-Aldrich, #C9754), GW4869 (Selleck Chemicals, #S7609), L778123 (MedChemExpress, #HY-16273A), LY294002 (Selleck Chemicals, #S1105), Y27632 (STEMCELL Technologies, #72304), and bafilomycin A1 (Sigma-Aldrich, #B1793-2UG) were dissolved in DMSO (Sigma-Aldrich, #D2650) and stored as aliquots in -20°C to minimize freeze–thaw cycles. We determined the maximum non-cytotoxic dose of each inhibitor by incubating 10,000 BFP-reporter HEK-293T cells with titrations of inhibitors for 48 hours in black 96-well plates (PerkinElmer, #6055660). Cell viability was determined using the Nano-Glo live cell assay according to the manufacturer’s instructions (Promega, #N2011). To determine the effect of the inhibitors on NANITE spread, we first transfected HEK-293T cells with either NANITE or Cas9 RNP plasmids in 6-well plates as above. The following day, 5,000 transfected HEK-293T cells were co-cultured with 5,000 BFP-reporter HEK-293T cells in the presence of inhibitors in 96-well plates. L778123, LY294002, Y27632, bafilomycin A1, colchicine, and DMSO were used at final concentrations of 1.6 µM, 3.1 µM, 25 µM, 1.25 nM, 1.6 µM, and 1 v/v% respectively. For GW4869 experiments, cells were transfected with NANITE plasmids targeting B2M rather than the BFP reporter, because GW4869 is fluorescent in the BFP channel. 5,000 transfected HEK-293T cells were co-cultured with 5,000 untransfected HEK-293T cells in the presence of 0.8 µM of GW4869. Flow cytometry was used to quantify editing and cell viability 48 hours after co-culture as described above. To simplify data interpretation, the fraction of edited cells in treated conditions were normalized to the fraction of edited cells in the untreated condition.

### Targeting NANITE

Anti-CD19 and anti-ACE2 single-chain variable fragments (scFvs) were designed as previously described^16,18,38^ and inserted upstream of VSVG in the NANITE plasmid with a P2A sequence. Mutations in the VSVG low-density lipoprotein receptor-binding region (K47Q, R354A) were introduced using appropriate primers and HiFi assembly. Plasmid sequences were verified and vesicles were produced as described above. Vesicles (12.5–50 μL) were incubated with 10,000 wild-type, CD19-expressing, or ACE2-expressing HEK-293T cells. Editing at the *B2M* locus was quantified 96 hours post-incubation by staining cells with 5 μL APC anti-human β2-microglobulin and 1 μL Zombie UV live-dead stain. B2M expression was quantified on an Attune NxT Flow Cytometer. The example flow gating strategy is shown in Fig. S3a, b.

For *in situ* experiments, NANITE plasmids were transfected into mCherry-expressing HEK-293T cells. The following day, transfected cells (5,000 cells) were mixed with 5,000 CD19-expressing HEK-293T cells (mNeonGreen) and 5,000 ACE2-expressing HEK-293T cells in a 96-well plate. B2M expression was quantified by flow cytometry as described above each day after co-culture. The gating strategy is shown in Fig. S3c. TGB610^41^, GFAP^42,63^, and LP1^43^ promoter sequences were purchased as gene fragments from Twist Bioscience. The CAG promoter in the NANITE plasmid was replaced with each of these promoters using HiFi assembly. Sequence-verified plasmids were transfected into BFP-reporter HEK-293T or HepG2 cells at 5 fmol and 24 fmol using PEI and jetPRIME reagents, respectively. An equimolar amount of plasmid was used to ensure cells received the same number of plasmids independent of the plasmid size. The following day, media were replaced to remove excess plasmid. BFP expression was quantified by flow cytometry 72 hours post-transfection as described above. To compare NANITE activity between cell lines, BFP expression was normalized to the CAG promoter condition.

### Administering NANITE in vivo

All mouse protocols were approved by the Institutional Animal Care and Use Committee (IACUC) at UCSF. Mice were housed in a barrier animal facility with standard 12-hour light/dark cycles at the Gladstone Institutes. All experiments were performed with 6–12-week-old female C57BL/6 mice (The Jackson Laboratory, #000664). Female mice were used exclusively to enable assessment of potential germline editing in ovaries, a key safety consideration for genome editing therapies with systemic exposure.

We initially packaged NANITE plasmids into lipid nanoparticles (LNPs) following previously published protocols^47^. Briefly, 50 mol% SM-102 (Cayman Chemical, #33474), 38.5 mol% cholesterol (MilliporeSigma, #C8667), 10 mol% DSPC (Avanti Research, #850365), and 1.5 mol% DMG-PEG2000 (Avanti Research, #880151) were mixed in anhydrous ethanol (Decon Laboratories, #V1016). The final lipid concentration of 12.5 mM. 9(10)-Nitrooleic acid (Cayman Chemical, #33896) was then added at 20 mol% of total lipid. Plasmids were diluted in 50 mM citrate buffer (pH 4) to a final concentration of 0.065 mg/mL. LNPs were formulated using the NanoAssemblr Ignite (Cytiva) with the following parameters: flow rate ratio of 1:3 (ethanol:aqueous), flow rate of 6 mL/min, and waste volumes of 0.15 mL and 0.05 mL at the beginning and end, respectively. LNPs were washed four times using 10 kDa Amicon Ultra centrifugal filters (MilliporeSigma, #UFC8010) using 1× PBS. Hydrodynamic diameters and polydispersity indices were measured using a BeNano 180 Zeta Max (Bettersize). Encapsulation efficiency was determined using Quant-iT PicoGreen assays (Thermo Fisher Scientific, #P7589) as previously described ^47^. For functional testing, LNPs (6–100 ng plasmid DNA) were incubated with BFP-reporter HEK-293T cells. Media were replaced the following day to remove excess LNPs. BFP and zsGreen expressions were quantified by flow cytometry as above. Following characterization, 0, 12.5, or 25 μg LNP-encapsulated plasmid was injected into C57BL/6 mice, matching the highest doses safely tested previously^47^. Transfection efficiency was assessed the following day, as plasmid expression peaks at approximately 24 hours post-transfection. Livers (either the left or right lateral lobe) were harvested and fixed for 24 hours in 4% buffered formaldehyde at 4°C, then transferred to 20 w/v% sucrose (Fisher Scientific, #BP220) in 1× PBS for 3 days. Livers were mounted in cryoblocks using Tissue-Plus O.C.T. Compound (Fisher Scientific, #23-730-571). Sections (20 μm) were cut using a Leica CM 3050 S cryostat, placed on microscope slides (Fisher Scientific, #22-037-246), and stored at −80°C. Sections were blocked with 1× Animal-Free Blocker (Vector Laboratories, #SP-5030) containing 0.03 v/v% Triton X-100 and 0.05 w/v% sodium azide (MilliporeSigma, #71289) for 1 hour at room temperature, then stained with DAPI (MilliporeSigma, #10236276001) for 30 minutes. Sections were coverslipped (Avantor, #48393-060) with Fluoromount-G (SouthernBiotech, #0100-01) and imaged using a Stellaris 5 confocal microscope (Leica) with identical settings across samples. No zsGreen fluorescence was detected, indicating no observable transfection.

We subsequently administered plasmids via hydrodynamic injection following UCSF standard protocols. For dose optimization studies, 15–34 pmol NANITE plasmid in saline (Teknova, #S5825) was injected at volumes equivalent to 10% of mouse body weight^48,50,51,64,65^. Transfection efficiency was assessed as above. The fraction of zsGreen-positive cells was quantified using CellProfiler by segmenting nuclei and zsGreen regions. Nuclei were masked with zsGreen regions to identify positive cells. The percentage of zsGreen-positive cells was calculated by dividing zsGreen-positive nuclei by total nuclei per field of view. For visualization, maximum and average intensity projections were applied to the DAPI and zsGreen channels, respectively, with identical thresholds across all images within an experiment.

To compare Cas9 RNP and NANITE plasmids *in vivo* for *Ttr* editing, both were administered at 20 pmol. Equimolar doses were used to ensure animals received equivalent plasmid copy numbers regardless of plasmid size. Body weights were recorded daily. Blood was collected from the submandibular vein at 1, 7, and 14 days post-injection using sterile lancets (Braintree Scientific, #GR 5MM) into capillary blood collection tubes (Sarstedt, #20.1280.100). Blood was kept on ice. To obtain serum, blood was centrifuged at 1,600 × g for 10 minutes at 4°C. Serum was collected and stored at −80°C until analysis. A sub-cohort of mice was euthanized 24 hours post-injection to quantify transfection, and remaining mice were euthanized at day 14. Tissues were harvested at each endpoint. All procedures were performed according to our IACUC-approved protocol.

To compare Cas9 RNP, NANITE (VSVG P127D) and NANITE plasmids *in vivo* for *Ttr* editing. We administered each plasmid at 20 pmol. Equimolar doses were used to ensure animals received equivalent plasmid copy numbers regardless of plasmid size. Blood was collected from the submandibular vein two weeks post-injection. Livers were harvested at the endpoint. All procedures were performed according to our IACUC-approved protocol.

### Quantifying NANITE, Cas9 and zsGreen expression in vivo

Transfection efficiency was quantified 24 hours post-injection by imaging, flow cytometry, and Western blotting of liver lysates. ZsGreen expression was quantified by imaging as outlined above. One liver section per mouse was analyzed. Flow cytometry on liver cells was performed following previously published protocols^66,67^. Fresh liver tissue (∼0.35 g) was minced using a sterile blade and digested with 50 U collagenase type I (Thermo Fisher Scientific, #17018029) in 1 mL Hank’s balanced salt solution (HBSS; Thermo Fisher Scientific, #14175145) supplemented with 5 mM CaCl (MilliporeSigma, #442909) at 37°C for 45 minutes with gentle shaking (300 rpm). The digested solution was filtered through a 70-μm cell strainer (Fisher Scientific, #22-363-548) and quenched with 2 mL HBSS supplemented with 2 v/v% FBS. Cell suspension (1 mL) was centrifuged at 500 × g for 5 minutes at 4°C to obtain a cell pellet. Hepatocytes were stained with 5 μL anti-CD95 antibody (BioLegend, #152620) and 1 μL Zombie NIR live-dead stain in HBSS for 30 minutes at 4°C. Cells were pelleted by centrifugation and washed with HBSS supplemented with 2 v/v% FBS. ZsGreen expression was quantified by flow cytometry on an Attune NxT Flow Cytometer. Gating is shown in Fig. S7a,b.

For Western blot analysis of Cas9 and NANITE expression in liver lysates, ∼20 mg liver tissue (from the left lateral, right lateral, and caudate lobes) was placed into 2 mL tubes prefilled with 1.4 mm ceramic beads (Omni International, #19-627) and 500 μL of RIPA lysis buffer with 1× protease inhibitors. Samples were kept on ice throughout processing. Tissue was homogenized using a BeadBug 6 (Benchmark Scientific) at 4,060 rpm for 9 seconds followed by 30 seconds of rest, repeated for 5 cycles. Lysates were centrifuged at >21,000 × g for 10 minutes at 4°C to remove debris. Supernatants were stored at −80°C until further processing. Samples were run on SDS-PAGE gels and transferred to 0.2 μm nitrocellulose membranes as described above. Membranes were blocked overnight at 4°C with 5 w/v% non-fat dry milk and 0.02 w/v% sodium azide (MilliporeSigma, #S2002-100G) in 1× TBS-T. Anti-FLAG antibody (MilliporeSigma, #F1804, 1:1,000) and anti-β-actin antibody (Cell Signaling, #4967, 1:1,000) in blocking solution were incubated with membranes overnight at 4°C. The anti-β-actin antibody was centrifuged at >19,000 × g for 10 minutes at 4°C to pellet aggregates prior to use. Membranes were washed 3–5 times for 5 minutes each with 1× TBS-T, then incubated with the appropriate secondary antibody (goat anti-mouse, Invitrogen, #62-6520, 1:1,000; or goat anti-rabbit, Invitrogen, #656120, 1:1,000) in blocking solution for 1 hour at room temperature. Membranes were washed 3–5 times and developed using chemiluminescent substrate (Thermo Fisher Scientific, #34580 or #34096). Images were acquired using a ChemiDoc MP imaging system. Raw Western blot images are shown in Fig. S10. Imaging and Western blot analyses were repeated identically for liver samples harvested at day 14 to assess persistence of expression. Raw Western blot images are shown in Fig. S10.

For quantification of plasmid DNA in liver genomic DNA, primers were designed to specifically amplify plasmid sequences at the ZsGreen coding region (Fw: ttcgccgaggacatcttgtc, Rv: cggggcaggagttcttgaag). Standard curves were generated using serial dilutions of Cas9 RNP or NANITE plasmids. Genomic DNA was quantified using a NanoDrop One spectrophotometer (Thermo Fisher Scientific, #ND-ONE-W) and normalized to 10 ng/µL prior to analysis. qPCR reactions were prepared in a final volume of 10 µL containing 5 µL of SYBR Green Universal Master Mix (Applied Biosystems, 4364344), 1 µL of primer mix (10 µM), 3 µL of nuclease-free water (Ambion, AM9937), and 1 µL of genomic DNA (10 ng total input). The cell number was calculated by assuming each diploid mouse cell had 7 pg of DNA.^68^ Reactions were assembled in 384-well optical plates (Applied Biosystems, A36931), sealed with optical adhesive film (Applied Biosystems, 4311971), and run on a QuantStudio 5 Real-Time PCR System (Applied Biosystems). Cycling conditions consisted of an initial incubation at 50°C for 2 min, followed by an initial denaturation step at 95°C for 10 min and 40 amplification cycles of 95°C for 15 s and 60°C for 1 min.

### Serum analyses

Circulating prealbumin (i.e, TTR) levels were quantified using the Mouse Prealbumin SimpleStep ELISA Kit (Abcam, #ab282297) according to the manufacturer’s instructions. Briefly, serum was diluted 1:100,000 in the kit dilution buffer. Samples or standards (50 μL) and 50 μL of antibody mix were added to assay plates and incubated for 1 hour at room temperature on an orbital shaker. Wells were washed three times with 300 μL 1× wash buffer, then given a final wash with 150 μL 1× wash buffer. TMB developing solution (100 μL) was added to each well, and plates were incubated for >15 minutes at room temperature to allow color development. The reaction was stopped by adding 100 μL of stop solution. Absorbance (λ = 450 nm) was measured immediately using a Spark 20M plate reader (Tecan).

We also quantified the serum sgRNA concentrations at 1 and 14 days post-injection. Serum samples were diluted 1:100 in 1× DNA/RNA Shield DirectDetect reagent (Zymo Research, #R1400). Standard curves were generated using unmodified *Ttr* sgRNA (Integrated DNA Technologies) in untreated mouse serum diluted 1:100 in 1× DNA/RNA Shield DirectDetect. Samples or standards (4 μL) were combined with 2.5 μL of Luna Universal qPCR Master Mix, 2 μL of reverse-transcription primer, 0.5 μL of probes (Custom TaqMan Small RNA Assay CTAACJP, Thermo Fisher Scientific), and 1 μL of nuclease-free water. Reactions were assembled in 384-well plates and sealed with optical adhesive film. RT-qPCR was run on a QuantStudio 5 Real-Time PCR System with the following settings: carryover prevention (25°C, 30 seconds), reverse transcription (55°C, 15 minutes), initial denaturation (95°C, 1 minute), and 45 cycles of denaturation (95°C, 10 seconds) and extension (60°C, 60 seconds) with a plate read. To test for neutralizing antibodies in the serum after treatment, we produced NANITE vesicles as described above and assessed their editing potency in the presence of treated mouse serum *in vitro*. We added 2 µL of serum to 10,000 BFP-reporter HEK-293T cells and 25 µL of vesicles in a total volume of 75 µL. Anti-VSVG antibodies at varying concentrations served as a positive control. After two days, editing was quantified by BFP expression using flow cytometry as described above.

To perform blood biochemistry, we collected serum at day 7 and day 14. Multiple blood collections within the first 7 days limited the volume available at day 7 per our animal protocol. We therefore only quantified alanine aminotransferase (ALT; Abcam #ab282882) and aspartate aminotransferase (AST; Abcam #ab325368) from day 7 samples by ELISA following the manufacturer’s protocol. Day 14 serum was submitted to the UC Davis Comparative Pathology Laboratory for Chem-11 panel analysis, which measured ALT, AST, alkaline phosphatase, total bilirubin, calcium, phosphorus, blood urea nitrogen, creatinine, glucose, total protein, and albumin. This timepoint was selected to capture cumulative effects. To enable comparison between timepoints, day 14 serum samples were also analyzed by ELISA, allowing us to generate a standard curve to convert ELISA values (ng/mL) to the activity units reported by the Chem-11 panel. Reference values for normal physiological ranges were taken from Charles River.

### Sequencing

Genomic DNA was extracted from fresh tissue using the Monarch Spin gDNA Extraction Kit (New England Biolabs, #T3010). Fresh liver tissue (∼20 mg from the left lateral, right lateral, and caudate lobes) or ovaries were placed into 200 μL tissue lysis buffer with 3 μL proteinase K. Tissues were minced in lysis buffer using sterile scissors, then incubated at 56°C with agitation (1,400 rpm) for 45 minutes. Tissue debris was removed by centrifugation at >12,000 × g for 3 minutes, and 3 μL RNase A was added to the supernatant. Samples were incubated for 5 minutes at 56°C with agitation (1,400 rpm). Genomic DNA binding and column purification were performed according to the manufacturer’s protocol.

Target regions were amplified by PCR using Q5 High-Fidelity DNA Polymerase (New England Biolabs, #M0492) and primers listed in Table S2. Potential off-target sites were predicted using Cas-OFFinder 2.4.1, prioritizing sites with NGG PAMs allowing up to three mismatches with one DNA or RNA bulge^69^. There were no predicted off-targets with less than three mismatches. Amplicons were column-purified using the QIAquick PCR Purification Kit (Qiagen, #28104) following the manufacturer’s protocol. Amplicon size and purity were verified by agarose gel electrophoresis, and concentration was determined using the Qubit dsDNA Quantification Assay (Thermo Fisher Scientific, #Q32854). Amplicons were submitted for Amplicon-EZ sequencing (∼50,000 reads per sample; Genewiz). Indel frequencies were quantified using CRISPResso2 with default parameters for an NHEJ experiment as previously published^70^.

For the loss-of-function experiments, sanger sequencing was used to quantify genome editing. Target regions were amplified by PCR using primers listed in Table S2. Amplicons were submitted to ELIM biopharmaceuticals, Inc for PCR cleanup and sequencing. Editing efficiencies were determined using the TIDE analysis webserver with default parameters as previously published^61^.

### *miniEDV* animal experiments

miniEDVs were produced as previously published^18^. Briefly, 28–30 million HEK-293T cells were seeded into T225 flasks (Thermo Fisher Scientific, #12-565-221) in 30 mL complete DMEM. The following day, cells were transfected with 15 µg pMD2.G, 30 µg TTR miniGag-Cas9, and 15 µg TTR miniGag-Pol plasmids using 181 µL Lipofectamine 3000 and 165 µL P3000 reagent (Thermo Fisher Scientific, #L3000015) in 10 mL of Opti-MEM I Reduced Serum Medium (Corning, #31-985-088). Prior to transfection, 15 mL of culture medium was removed from each flask. Six hours post-transfection, the medium was replaced with Opti-MEM I supplemented with 10% FBS (Avantor, #97068-085) and 1x Viral Boost reagent (AllStem, #VB100). Particles were harvested 48 hours post-transfection by centrifugation at 2,000 x g for 10 min to remove cellular debris. The supernatant was filtered through a 0.45-µm PES Rapid-Flow sterile filter unit (Thermo Fisher Scientific, #165-0045) and transferred to Thinwall Ultra-Clear tubes (Beckman Coulter, #NC9146666). A sucrose cushion of 50 mM Tris-HCl (pH 7.4), 100 mM NaCl, 0.5 mM EDTA, and 20 w/v% sucrose was layered beneath the supernatant. Samples were ultracentrifuged at 100,000 x g for 1.5 hr at 4°C using an SW 32 Ti swinging-bucket rotor (Beckman Coulter, #5043-30-2260) in an Optima XE-90 ultracentrifuge (Beckman Coulter). Pellets were resuspended in 25–50 µL cold PBS and stored at −80°C. Particles were titered by RT-qPCR and nanoFCM as described above. All animal procedures were conducted in accordance with the approved IACUC protocol. miniEDVs were administered by tail vein injection at UCSF at doses of 0, 3.2 × 10^10^ or 5.5 × 10^10^ particles per mouse. Serum, body weight, and editing analyses were performed as described above.

### Liver histopathology

Liver lobes were harvested and fixed in 5 mL 4% buffered formaldehyde at 4°C for 24 hours. Following fixation, tissues were transferred to 70% (v/v) ethanol and stored at 4°C until submission to the Gladstone Histology and Light Microscopy Core for processing. Liver lobes were placed in cassettes, processed, and embedded in paraffin. Sections were mounted onto glass slides and stained with hematoxylin and eosin (H&E). Slides were imaged using an Axioscan 7 slide scanner (Zeiss) at 20× magnification. Whole liver sections were scored for signs of hepatotoxicity and fused multinucleated cells by a licensed pathologist in a blinded manner (Histowiz). One section was scored per mouse.

### Software and Statistics

Flow cytometry data were analyzed using FlowJo 10.10.0 as indicated. Images were processed for visualization in (Fiji Is Just) ImageJ 2.16.0/1.54p. Image analyses were performed in CellProfiler 4.2.8 or (Fiji Is Just) ImageJ 2.16.0/1.54p as indicated. Cas-OFFinder 2.4.1 was used to predict off-target sites for genome editing. CRISPResso2 was used to analyze sequencing data and determine editing efficiencies. Data were plotted in GraphPad Prism v.10.6.1 or Microsoft Excel v.16.104. Statistical significance was determined using GraphPad Prism v.10.6.1 as indicated. Figures were created using Adobe Illustrator 28.4.1.

## Data availability

Plasmids are available from Addgene (#258772 - 258781). All other data and materials are available from the authors upon request.

## Code availability

Bespoke code was not used in this work.

## Acknowledgements

We would like to thank Kevin Pastores from the Gladstone Flow Cytometry Facility for help with flow cytometry. The Gladstone Flow Cytometry core is funded by NIH S10 RR028962 and the James B. Pendleton Charitable Trust. We would also like to thank Alex Liu from the Gladstone Light and Microscopy Core. We are grateful to Alex Marson for initial help with animal experiments. We thank all members of the Doudna laboratory for their thoughtful input on this manuscript. We acknowledge the use of Claude Opus 4.8 for text editing. Funding was provided by National Heart, Lung, and Blood Institute grant 1R21HL173710-01, Lawrence Livermore National Labs PROTECT grant, DE-AC52-07NA27344, the Gladstone Institutes and the Howard Hughes Medical Institute. J.A.D also receives support from NIH/NIAID (U54AI170792, U19AI135990, UH3AI150552 and U01AI142817), NIH/NINDS (U19NS132303), NIH/NHLBI (R21HL173710), NSF (2334028), DOE (DE-AC02-05CH11231, 2553571 and B656358); Lawrence Livermore National Laboratory, Apple Tree Partners (24180), UCB-Hampton University Summer Program, Mr. Li Ka Shing, Koret-Berkeley-TAU, Emerson Collective and the Innovative Genomics Institute (IGI). J.L.Y.W was founded by a Natural Sciences and Engineering Research Council of Canada Postdoctoral Fellowship. Mason T. Hooks was funded by a NIH T32 training grant. We would also like to acknowledge the generous support of the James B. Pendleton Charitable trust.

## Author information

***Innovative Genomics Institute; University of California, Berkeley; Berkeley CA, USA*** Wayne Ngo, Daniel Rosas-Rivera, Kevin M. Wasko, Min Hyung Kang, Shaan Gogna, Jingkun Zeng, Mason T. Hooks, Jamie L.Y. Wu, Jennifer A. Doudna

***Gladstone Institute of Data Science and Biotechnology; San Francisco, CA, USA***

Wayne Ngo, Daniel Rosas-Rivera, Jingkun Zeng, Jamie L.Y. Wu, Jennifer A. Doudna

***California Institute for Quantitative Biosciences, University of California, Berkeley; Berkeley, CA, USA***

Wayne Ngo, Jingkun Zen, Jamie L.Y. Wu, Jennifer A. Doudna

***Department of Molecular and Cell Biology, University of California, Berkeley; Berkeley, CA, USA***

Kevin M. Wasko, Min Hyung Kang, Mason T. Hooks, Jennifer A. Doudna

***Department of Medicine, University of California, San Francisco; San Francisco, CA, USA***

Longhui Qiu

***Gladstone-UCSF Institute of Genomic Immunology; San Francisco, CA, USA***

Zhongmei Li, Jennifer A. Doudna

***Howard Hughes Medical Institute, University of California, Berkeley; Berkeley CA, USA***

Jennifer A. Doudna

***Molecular Biophysics and Integrated Bioimaging Division, Lawrence Berkeley National Laboratory; Berkeley, CA, USA***

Jennifer A. Doudna

***Department of Chemistry, University of California, Berkeley; Berkeley, CA, USA***

Jennifer A. Doudna

***Li Ka Shing Center for Genomic Engineering, University of California, Berkeley, Berkeley, CA, USA***

Jennifer A. Doudna

## Contributions

W.N. and J.A.D conceptualized the study. WN. and J.A.D developed the methodology. All authors carried out the investigation. W.N., D.R.R, and J.A.D created visual representations.

W.N and J.A.D wrote the initial draft with substantial contributions from all authors. All authors reviewed and edited the manuscript. W.N. and J.A.D were responsible for funding acquisition and project administration.

***Corresponding author***

Correspondence to <u>Jennnifer A. Doudna</u>

## Additional information

***Publisher’s note*** Springer Nature remains neutral with regard to jurisdictional claims in published maps and institutional affiliations

## Supplementary information

### Supplementary Information

Supplementary Figs. 1 - 10 and Supplementary Table 1 - 2.

