## Supplementary Figures 1 - 10; Supplementary Table 1 - 2 for "Amplified genome editing by *in vivo* editor production"

**Table of Contents**

Supplementary Fig. 1. Design and performance of the BFP reporter.	3

Supplementary Fig. 2. Effect of promoter-to-IRES distance on zsGreen expression.	4

Supplementary Fig. 3. Flow gating strategy for the specificity experiment.	6

Supplementary Fig. 4. Transfection efficiency of producer cells.	7

Supplementary Fig. 5. NANITE plasmids can be packaged into DNA-LNPs, but failed to transfect liver cells in vivo.	8

Supplementary Fig. 6. Optimization of plasmid dose for hydrodynamic injection.	9

Supplementary Fig. 7. ZsGreen expression in liver hepatocytes after hydrodynamic injections.	10

Supplementary Fig. 8. Benchmarking NANITE genome editing to miniEDV genome editing.	11

Supplementary Fig. 9. Humoral response to NANITE-plasmid treated or miniEDV-treated mice.	12

Supplementary Fig. 10. Unprocessed Western blot images.	13

Supplementary Table 1. Cas9 spacer sequences.	14

Supplementary Table 2. Sequencing primers	14

References	15

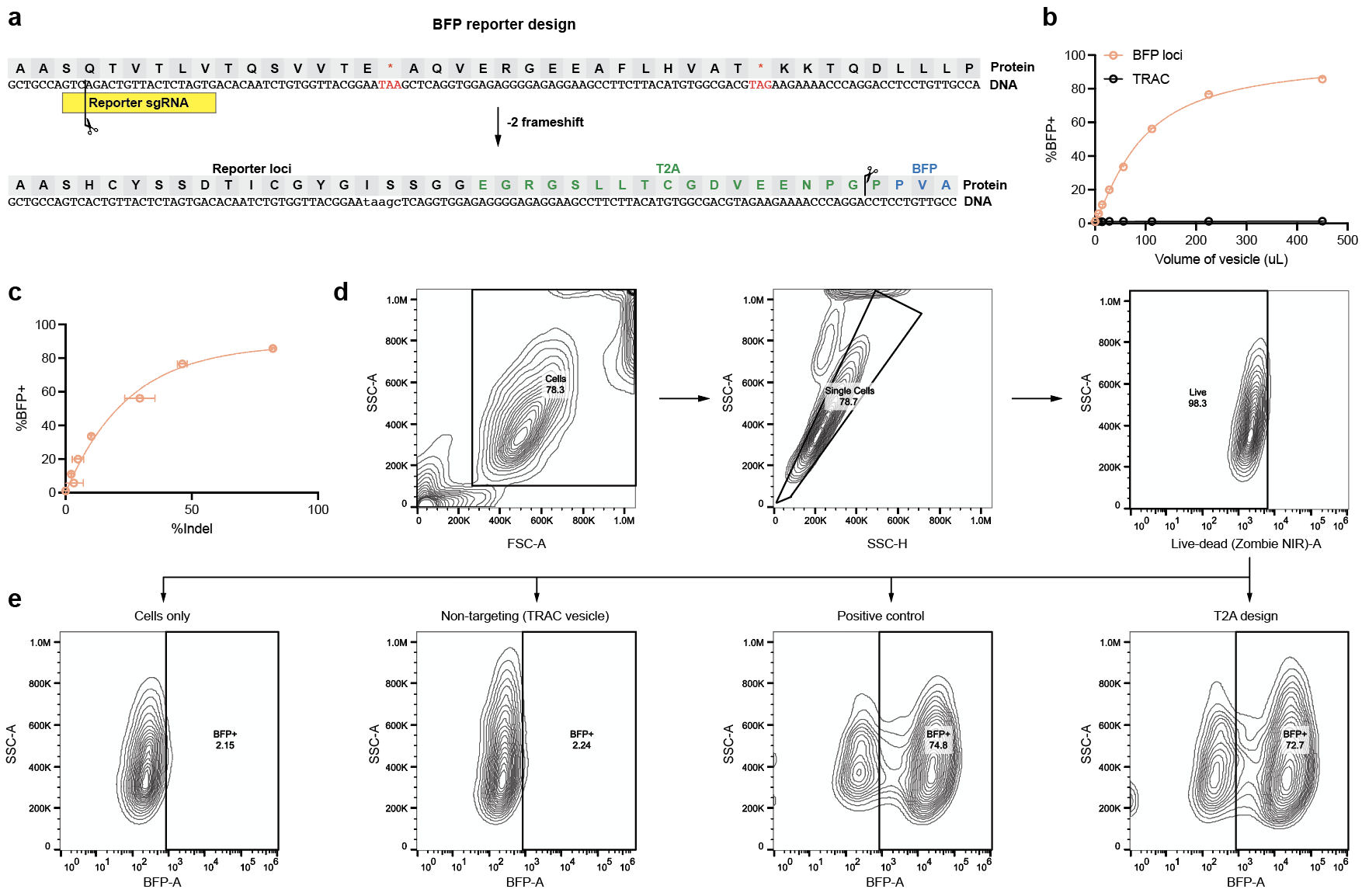

### Supplementary Fig. 1. Design and performance of the BFP reporter.

**(a)** A +2 frameshift is introduced into the reporter locus upstream of a T2A-BFP sequence, resulting in multiple premature stop codons that prevent BFP translation. Indels generated by genome editing may correct the frameshift and restore BFP expression. Multiple copies of the reporter were integrated into cells via lentiviral transduction. **(b)** Percentage of BFP+ cells after incubation with miniEDVs targeting either the BFP reporter or the negative control *TRAC* locus. BFP expression is specific to editing at the BFP locus. **(c)** Comparison of BFP+ cell percentage to indel frequency at the reporter locus following miniEDV incubation. BFP expression is proportional to editing at the reporter locus but plateaus at 60–80% BFP+. Data are presented as mean ± SD of three technical replicates. **(d,e)** Flow cytometry gating strategy used to determine the percentage of BFP+ cells. **(e)** Representative contour plots showing BFP expression in untreated, non-targeting control, positive control, and T2A design conditions.

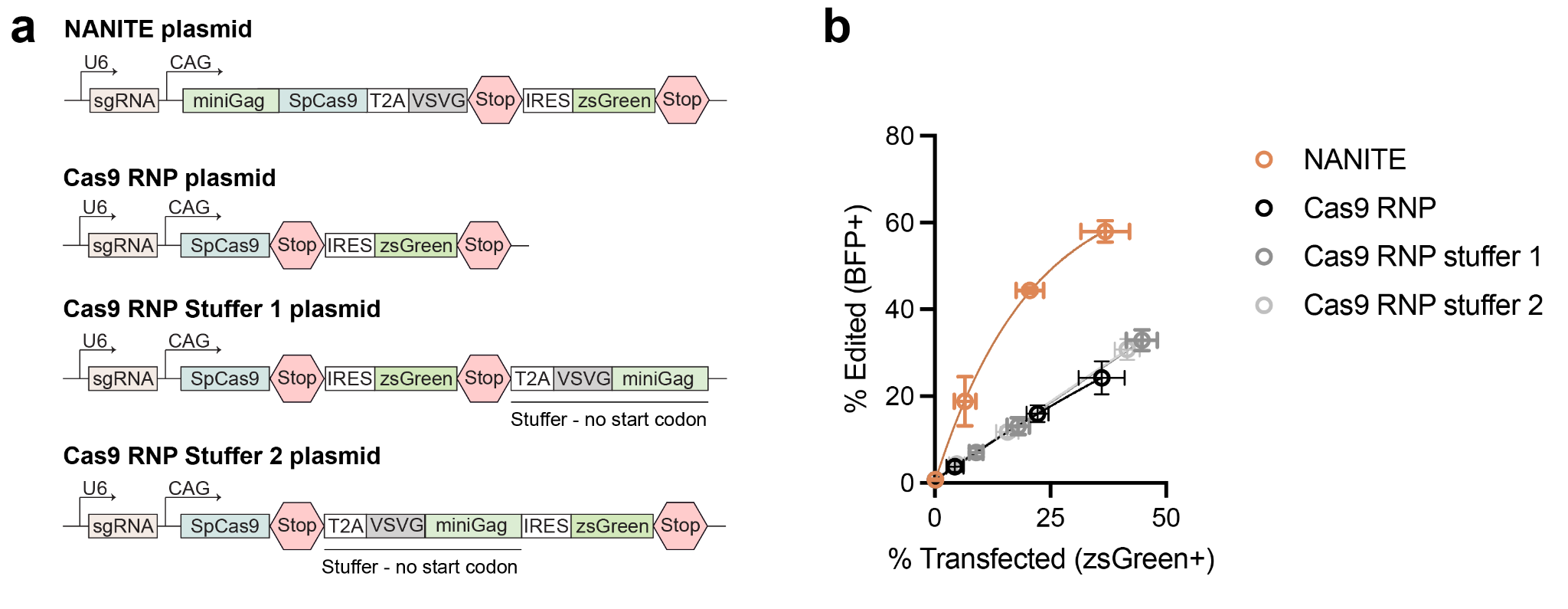

### Supplementary Fig. 2. Effect of promoter-to-IRES distance on zsGreen expression.

**(a)** Schematics of plasmid designs. An inert stuffer sequence (T2A-VSVG-miniGAG lacking a start codon) was inserted in the Cas9 RNP plasmid in different positions. Schematics are not to scale. **(b)** Fraction of edited cells relative to transfected cells following transfection with NANITE- or Cas9 RNP-plasmids with stuffers. The Cas9 RNP-plasmid behaved similarly with or without stuffers. Data are presented as mean ± SD of three replicates.

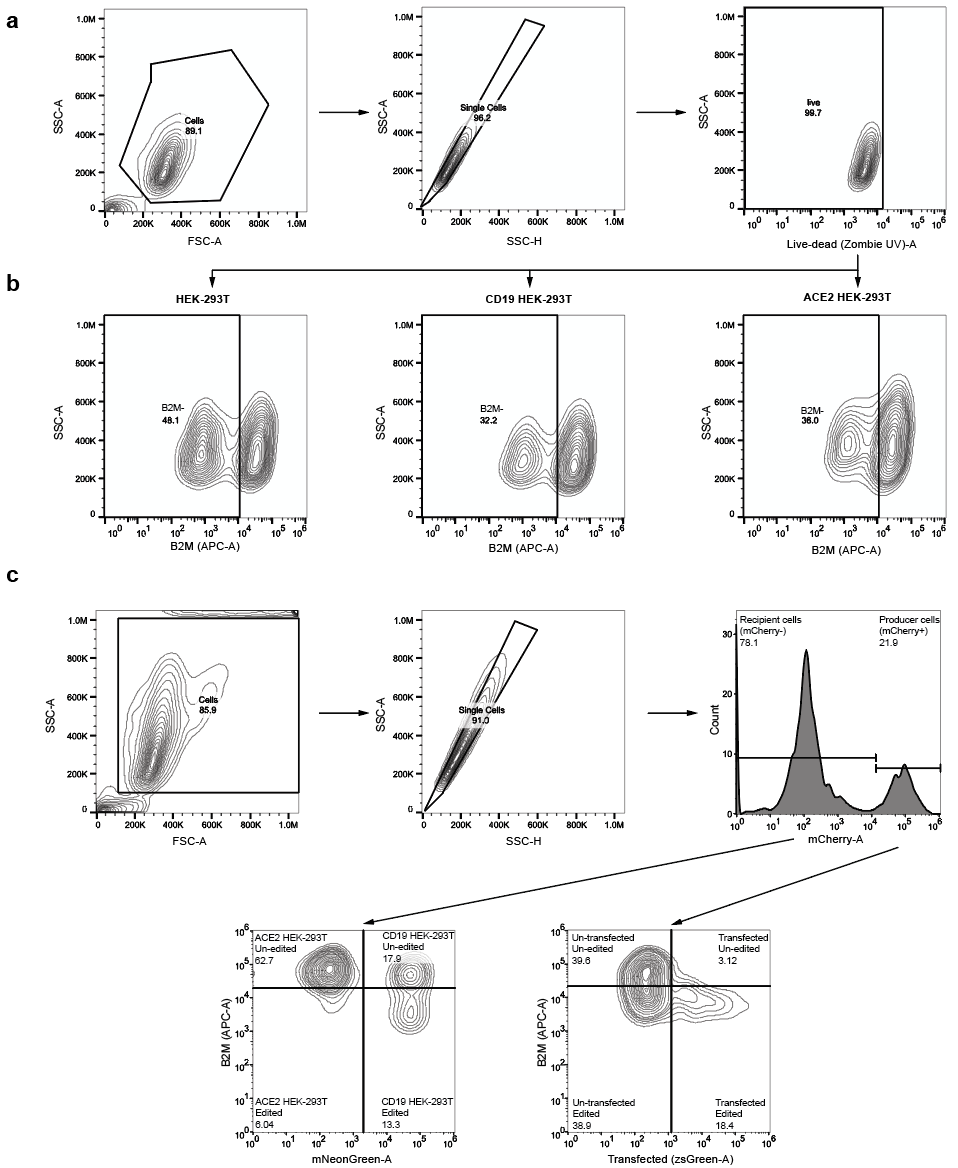

### Supplementary Fig. 3. Flow gating strategy for the specificity experiment.

**(a,b)** Flow gating strategy for data in Figure 3C. **(b)** Representative contour plots showing HEK-293T, CD19-expressing HEK-293T, or ACE2-expressing HEK-293T cells incubated with 25 μL VSVG-vesicles. **(c)** Flow gating strategy for experiment in Figure 3E.

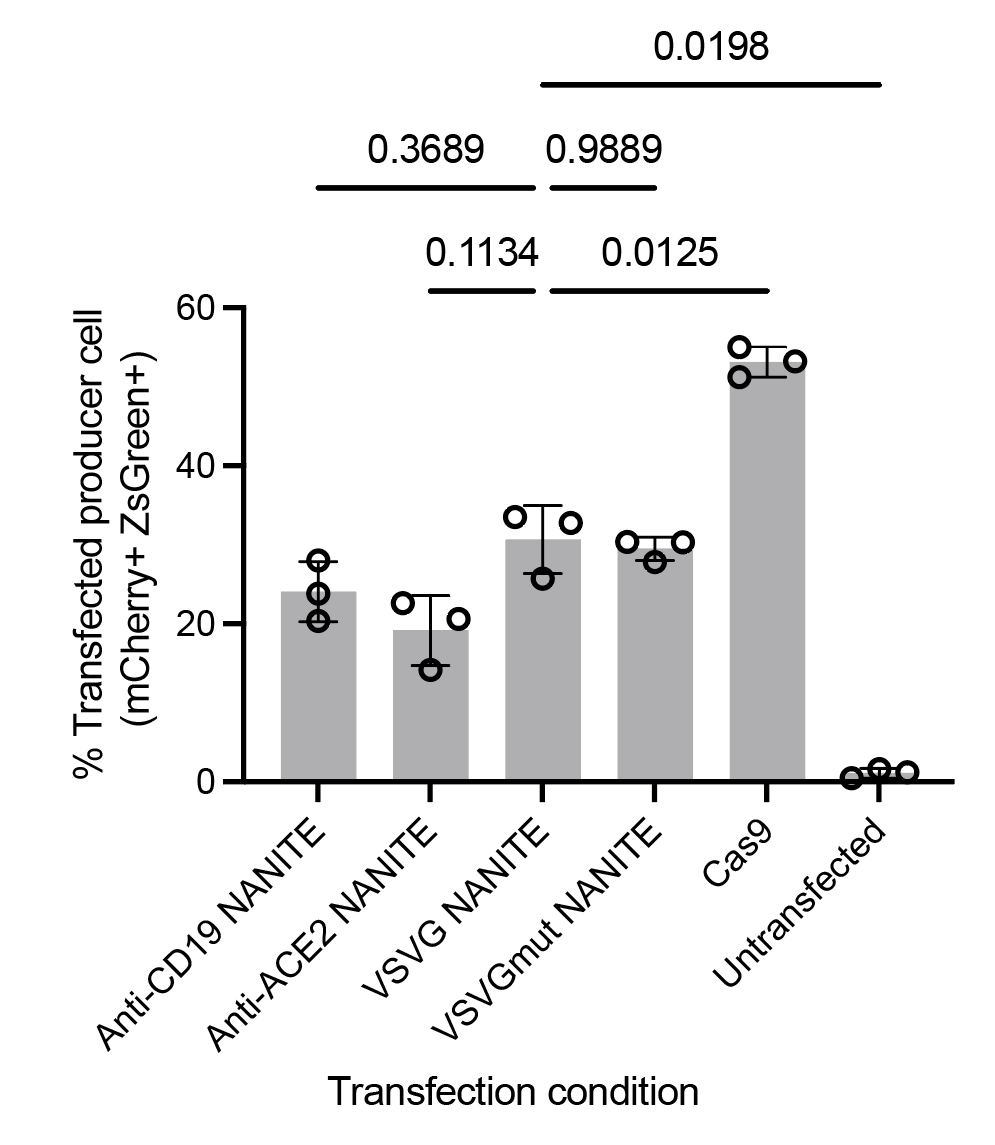

### Supplementary Fig. 4. Transfection efficiency of producer cells.

Percentage of transfected producer cells one day after transfection with 2500 ng of the indicated plasmid. NANITE conditions showed similar transfection efficiencies. The Cas9 RNP-plasmid condition had a higher transfection efficiency, because it had a lower molecular weight, so normalizing by mass meant a higher molar amount was transfected. Statistical differences were determined by Welch's ANOVA test with Dunnett's T3 multiple comparison test. p-values are as indicated. Data are presented as mean ± SD of three replicates.

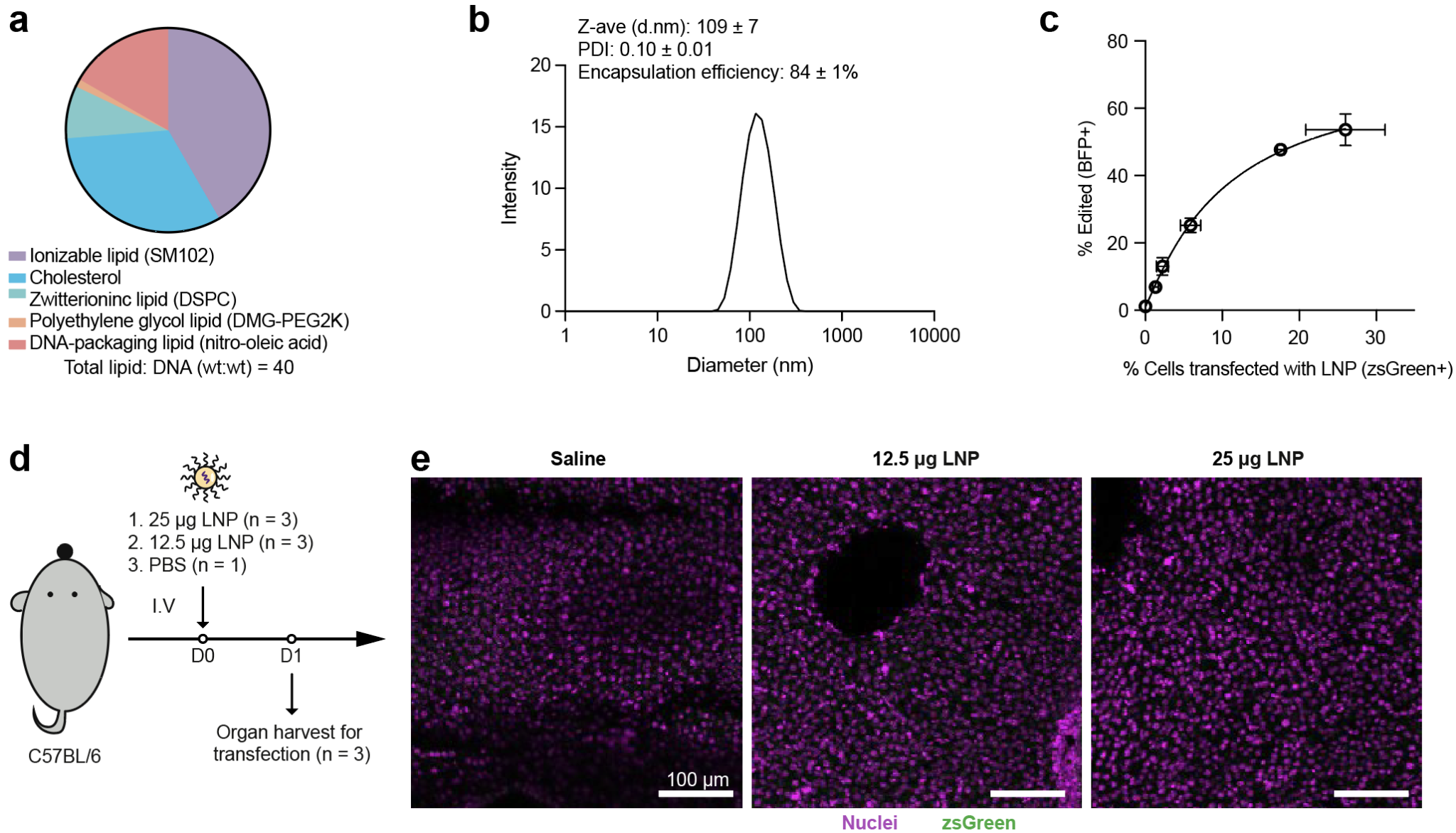

### Supplementary Fig. 5. NANITE plasmids can be packaged into DNA-LNPs, but failed to transfect liver cells *in vivo*.

**(a)** Lipid composition of DNA-LNPs. **(b)** Size distribution of DNA-LNPs after purification. The average hydrodynamic diameter, polydispersity index (PDI), and encapsulation efficiency are indicated. Data are presented as mean ± SD of three independent batches. **(c)** Percentage of edited relative to transfected BFP-reporter cells following DNA-LNP transfection. More edited cells than transfected cells were observed as expected. Data are presented as mean ± SD of three biological replicates. **(d)** Schematic of *in vivo* experiments. Mice were injected with DNA-LNPs at the highest doses reported in the literature[^1^](https://app.readcube.com/library/263feb6a-21a9-473a-9afe-b8966a8b0638/all?uuid=3460070843341212&item_ids=263feb6a-21a9-473a-9afe-b8966a8b0638:ff1d09cf-ad50-4414-997d-254ab395ad57). Organs were harvested 24 hours post-injection to assess liver transfection by zsGreen expression. **(e)** Representative immunofluorescence images of liver sections showing nuclei (magenta) and zsGreen (green). No zsGreen was detected. Scale bar as indicated.

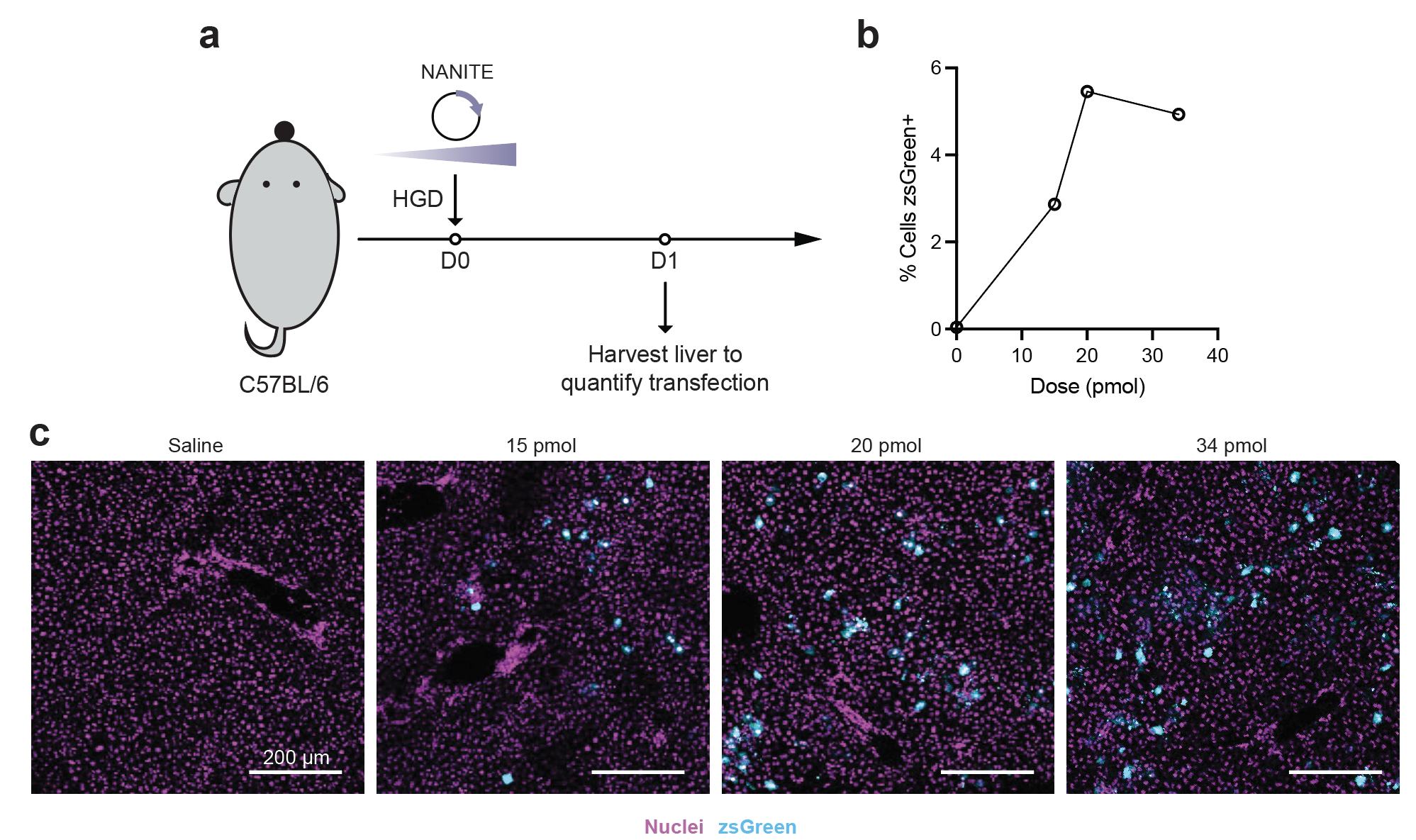

### Supplementary Fig. 6. Optimization of plasmid dose for hydrodynamic injection.

**(a)** Schematic of mouse experiments. NANITE plasmid was hydrodynamically injected (HGD) into mice at increasing doses. Livers were harvested 24 hours post-injection to assess transfection by zsGreen expression. **(b)** Percentage of liver cells expressing zsGreen, quantified using CellProfiler from images shown in **(c)**. Transfection plateaued at approximately 5% of liver cells. **(c)** Representative immunofluorescence images of liver sections. Nuclei are stained with DAPI (magenta). zsGreen is shown in cyan. One mouse per dose was used. Scale bar as indicated.

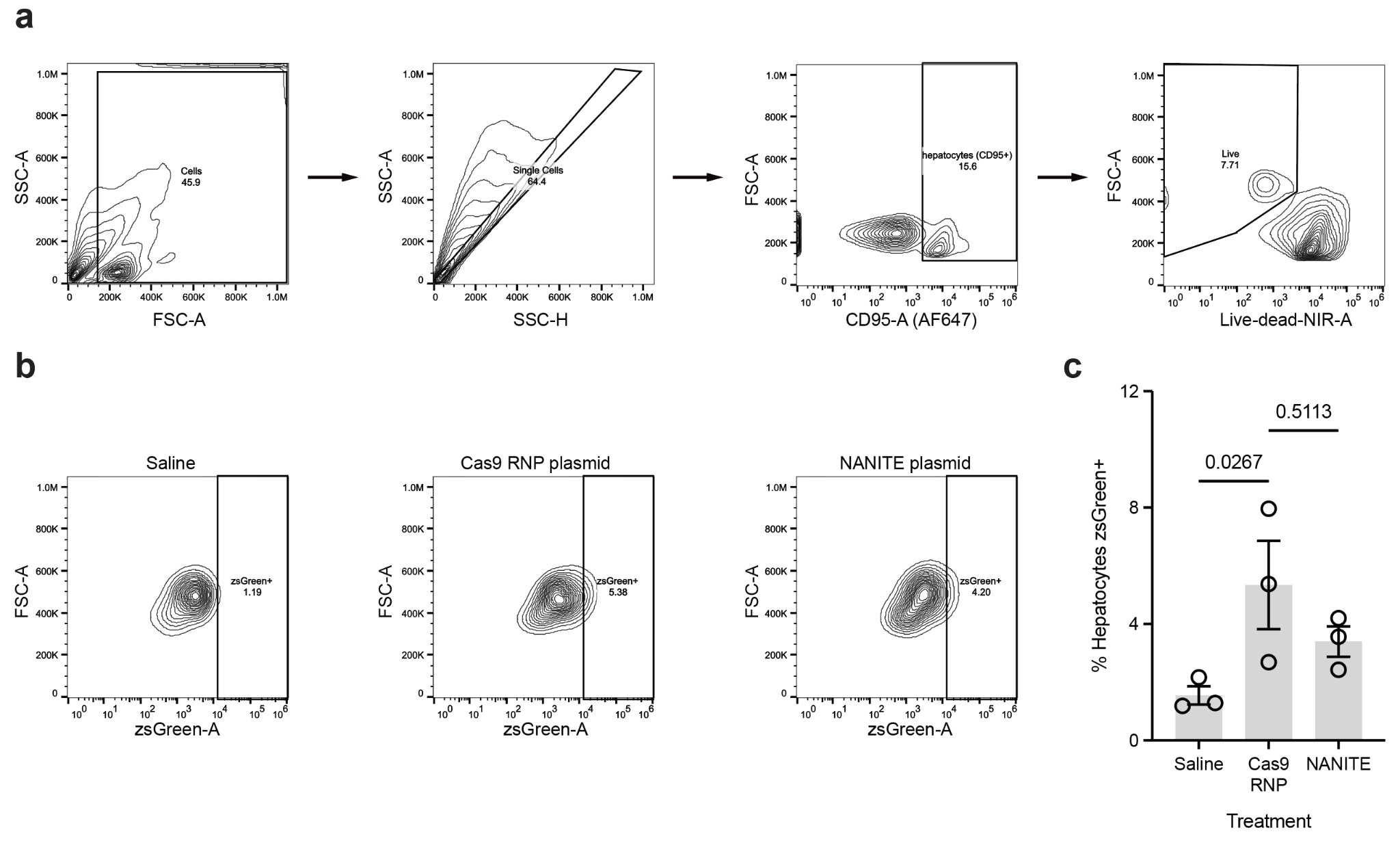

### Supplementary Fig. 7. ZsGreen expression in liver hepatocytes after hydrodynamic injections.

**(a,b)** Flow gating strategy to detect zsGreen expression in CD95+ hepatocytes isolated from mouse livers. **(b)** Representative contour plots showing zsGreen expression in mice injected with saline, Cas9 RNP-plasmid, or NANITE-plasmid. **(c)** Percentage of zsGreen+ hepatocytes quantified by flow cytometry. Data are presented as the mean ± SEM of three mice. Statistical significance was determined by one-way ANOVA on log-transformed data with Tukey's multiple comparison test. No significant difference in the proportion of transfected cells was observed between Cas9 RNP and NANITE plasmid groups.

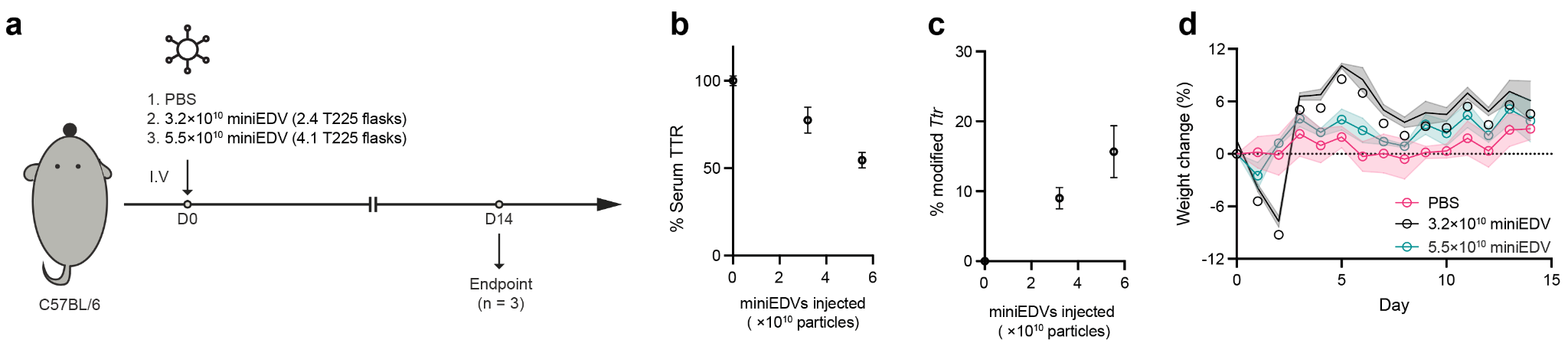

### Supplementary Fig. 8. Benchmarking NANITE genome editing to miniEDV genome editing.

**(a)** Mice were administered different doses of miniEDVs. Serum TTR and genome editing was quantified on day 14. **(b)** Serum TTR levels were quantified by ELISA on day 14, showing a dose-dependent reduction with increasing miniEDV dose. **(c)** Editing at the *Ttr* locus on day 14. At 20 pmol, NANITE achieved editing comparable to 5.5 × 10^10^ miniEDV particles per mouse. **(d)** Body weight over time. Mice treated with the high dose of miniEDVs exhibited transient weight loss one day after injection. Data are the mean ± SEM of three mice.

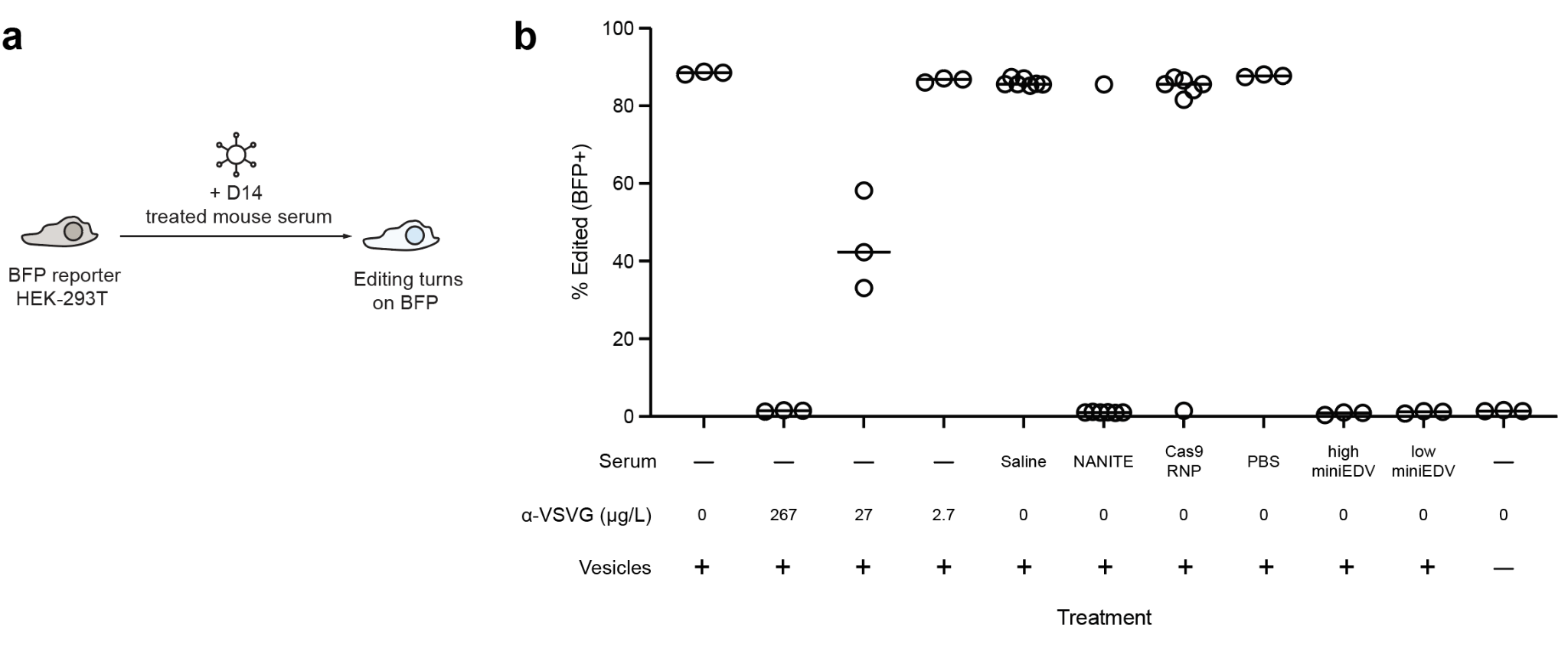

### Supplementary Fig. 9. Humoral response to NANITE-plasmid treated or miniEDV-treated mice.

**(a)** BFP reporter cells were incubated with NANITE-encoded vesicles after pre-incubation with the indicated mice serums or antibody positive controls. Editing was quantified by flow cytometry after two days after incubation. **(b)** Fraction of edited cells. NANITE-plasmid treatment elicits a similar level of neutralizing antibodies as miniEDVs injected at low (3.2 × 10^10^ particles) or high dose (5.5 × 10^10^ particles). The mean of each group is shown. Each point represents one biological replicate.

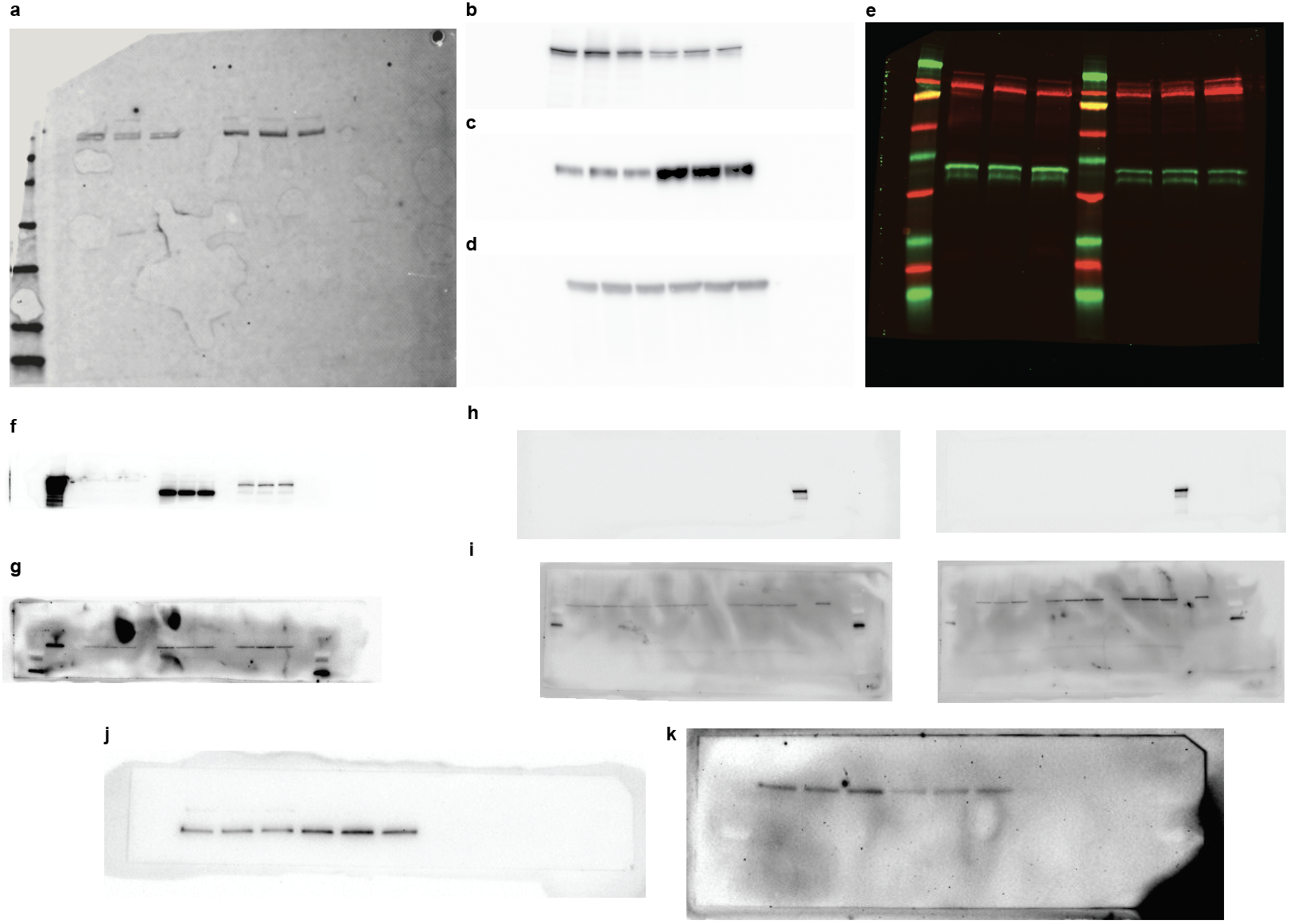

### Supplementary Fig. 10. Unprocessed Western blot images.

**(a)** Unprocessed Western blot for Fig. 1F. **(b–d)** Unprocessed Western blots for Extended Data Fig.1a . **(e)** Unprocessed Western blot for Extended Data Fig. 1b **(f,g)** Unprocessed Western blots for Extended Data Fig. 6a. **(h,i)** Unprocessed Western blots for Extended Data Fig. 6g. (**j,k)** Unprocessed Western blots for Fig. 5c

| Supplementary Table 1. Cas9 spacer sequences. | | |
| --- | --- | --- |
| **Name** | **Spacer sequence** | **PAM** |
| BFP reporter | CACTAGAGTAACAGTCTGAC | TGG |
| B2M | GAGTAGCGCGAGCACAGCTA | AGG |
| Ttr | TTACAGCCACGTCTACAGCA | GGG |

| Supplementary Table 2. Sequencing primers | | |
| --- | --- | --- |
| **Name** | **Type of Sequencing** | **Sequence** |
| BFPreporter.rev | Sanger | GTCCGCCGGGTACAAAGTTTC |
| BFPreporter.for | Sanger | AGCGGAAAGATGGCCGCTTC |
| Sanger-TTR.fwd | Sanger | ATGTTATCATAATGGGATCAGCATG |
| Sanger-TTR.rev | Sanger | CTCTCTCTGAGCCCTCTAGCTG |
| TTR.for | NGS | ACACTCTTTCCCTACACGACGCTCTTCCGATCTACATTTCTCTTGTCTCCTCTG |
| TTR.rev | NGS | GACTGGAGTTCAGACGTGTGCTCTTCCGATCTAACCTGGAACCGGTCCCATG |
| OT1.for | NGS | ACACTCTTTCCCTACACGACGCTCTTCCGATCTAAGGACACGCTCTGGGGCTG |
| OT1.rev | NGS | GACTGGAGTTCAGACGTGTGCTCTTCCGATCTCATGGAAGCCGTATTCAACC |
| OT2.for | NGS | ACACTCTTTCCCTACACGACGCTCTTCCGATCTTCAAGGGAGGTTGTATGGGAG |
| OT2.rev | NGS | GACTGGAGTTCAGACGTGTGCTCTTCCGATCTTTCTGCTAAATGGACATAGTAT |
| OT3.for | NGS | ACACTCTTTCCCTACACGACGCTCTTCCGATCTACACGCTCTGGGGCTGCTGG |
| OT3.rev | NGS | GACTGGAGTTCAGACGTGTGCTCTTCCGATCTCATGGAAGCCGTATTCAACC |
| OT4.for | NGS | ACACTCTTTCCCTACACGACGCTCTTCCGATCTTGCCCAAGTTTCACTTTTCT |
| OT4.rev | NGS | GACTGGAGTTCAGACGTGTGCTCTTCCGATCTGGACATGGAAACACAGACGG |
| OT5.for | NGS | ACACTCTTTCCCTACACGACGCTCTTCCGATCTTCCCGCTGCGCCTCTCTCTA |
| OT5.rev | NGS | GACTGGAGTTCAGACGTGTGCTCTTCCGATCTTCACTTGACCCCCACTCCCT |
| OT6.for | NGS | ACACTCTTTCCCTACACGACGCTCTTCCGATCTTGTATGAAATTAGTAGCACATTAT |
| OT6.rev | NGS | GACTGGAGTTCAGACGTGTGCTCTTCCGATCTGACAGACTCCAGGACATGAA |
| OT7.for | NGS | ACACTCTTTCCCTACACGACGCTCTTCCGATCTCTAACCCTGCCTCTGGGCAA |
| OT7.rev | NGS | GACTGGAGTTCAGACGTGTGCTCTTCCGATCTCCCCAATAGAGTTTGACAGTTCT |
| OT8.for | NGS | ACACTCTTTCCCTACACGACGCTCTTCCGATCTGAGCCACTTAAAAGAGATTC |
| OT8.rev | NGS | GACTGGAGTTCAGACGTGTGCTCTTCCGATCTGTTCCTCCTATGGGCTGCA |
| OT9.for | NGS | ACACTCTTTCCCTACACGACGCTCTTCCGATCTGCACTGCTTACATTGGCTTC |
| OT9.rev | NGS | GACTGGAGTTCAGACGTGTGCTCTTCCGATCTATCACGCTCACATGCAGCT |
| OT10.for | NGS | ACACTCTTTCCCTACACGACGCTCTTCCGATCTAGAGGTGGGGAGGAGGAGAG |
| OT10.rev | NGS | GACTGGAGTTCAGACGTGTGCTCTTCCGATCTGGAGAGCCTATCCTGAGCCA |
